# The Repertoire of Mutational Signatures in Human Cancer

**DOI:** 10.1101/322859

**Authors:** Ludmil B Alexandrov, Jaegil Kim, Nicholas J Haradhvala, Mi Ni Huang, Alvin WT Ng, Yang Wu, Arnoud Boot, Kyle R Covington, Dmitry A Gordenin, Erik N Bergstrom, S M Ashiqul Islam, Nuria Lopez-Bigas, Leszek J Klimczak, John R McPherson, Sandro Morganella, Radhakrishnan Sabarinathan, David A Wheeler, Ville Mustonen, the PCAWG Mutational Signatures Working Group, Gad Getz, Steven G Rozen, Michael R Stratton

**Affiliations:** Department of Cellular and Molecular Medicine and Department of Bioengineering and Moores Cancer Center, University of California, La Jolla, San Diego, CA, USA; Broad Institute, Cambridge, MA, USA; Center for Cancer Research, Massachusetts General Hospital, Boston, MA 02129, USA; Centre for Computational Biology and Programme in Cancer & Stem Cell Biology, Duke NUS Medical School, Singapore; Human Genome Sequencing Center, and Dan L. Duncan Cancer Center, Baylor College of Medicine, Houston, TX, USA; Genome Integrity and Structural Biology Laboratory, National Institute of Environmental Health Sciences, US National Institutes of Health, Research Triangle Park, NC, USA; Institute for Research in Biomedicine (IRB Barcelona), The Barcelona Institute of Science and Technology, Baldiri Reixac, 10, 08028 Barcelona, Spain; Research Program on Biomedical Informatics, Universitat Pompeu Fabra, Barcelona, Spain; Institució Catalana de Recerca i Estudis Avançats (ICREA), Barcelona, Spain; Integrative Bioinformatics Support Group, National Institute of Environmental Health Sciences, US National Institutes of Health, Research Triangle Park, NC, USA; Wellcome Sanger Institute, Wellcome Genome Campus, Hinxton, Cambridge CB10 1SA, UK; Organismal and Evolutionary Biology Research Programme, Department of Computer Science, Institute of Biotechnology, University of Helsinki, Helsinki, Finland; Department of Pathology, Massachusetts General Hospital, Boston, MA 02114, USA; Harvard Medical School, Boston, MA 02215, USA

## Abstract

Somatic mutations in cancer genomes are caused by multiple mutational processes each of which generates a characteristic mutational signature. Using 84,729,690 somatic mutations from 4,645 whole cancer genome and 19,184 exome sequences encompassing most cancer types we characterised 49 single base substitution, 11 doublet base substitution, four clustered base substitution, and 17 small insertion and deletion mutational signatures. The substantial dataset size compared to previous analyses enabled discovery of new signatures, separation of overlapping signatures and decomposition of signatures into components that may represent associated, but distinct, DNA damage, repair and/or replication mechanisms. Estimation of the contribution of each signature to the mutational catalogues of individual cancer genomes revealed associations with exogenous and endogenous exposures and defective DNA maintenance processes. However, many signatures are of unknown cause. This analysis provides a systematic perspective on the repertoire of mutational processes contributing to the development of human cancer including a comprehensive reference set of mutational signatures in human cancer.

## INTRODUCTION

Somatic mutations in cancer genomes are caused by mutational processes of both exogenous and endogenous origins that have operated during the cell lineage between the fertilised egg and the cancer cell^1^. Each mutational process may involve components of DNA damage/modification, DNA repair and DNA replication, any of which may be normal or abnormal, and generates a characteristic mutational signature that may incorporate base substitutions, small insertions and deletions, genome rearrangements, and chromosome copy number changes^2^. The catalogue of mutations from an individual cancer genome may have been generated by multiple mutational processes and thus incorporates multiple superimposed mutational signatures. Therefore, in order to systematically characterise the mutational processes contributing to cancer, mathematical methods have been developed that can be used to *(i)* decipher mutational signatures from a set of somatic mutational catalogues, *(ii)* estimate the numbers of mutations attributable to each signature in each sample, and *(iii)* annotate each mutation class in each tumour with the probability of arising from each signature^3–15^.

Previous studies of multiple cancer types identified >30 single base substitution signatures, some of known but many of unknown aetiologies, some ubiquitous and others rare, some part of normal cell biology and others associated with abnormal exposures or operative during neoplastic progression^6,16–27^. Six genome rearrangement signatures have also been identified in breast cancer^18^ and further patterns of rearrangements have been described^13,28–30^. However, analysis of other mutation classes has been relatively limited^31,17,18,32,33^.

Thus far, mutational signature analysis has predominantly used cancer exome sequences. However, the many fold greater numbers of somatic mutations in whole-genome sequences provide substantially increased power for signature decomposition, enabling better separation of partially correlated signatures and extraction of signatures that contribute relatively small numbers of mutations. Furthermore, technical artefacts and differences in sequencing technologies and mutation calling algorithms can themselves generate mutational signatures. Therefore, the uniformly processed and highly curated sets of all classes of somatic mutations from the 2,780 cancer genome sequences of the Pan Cancer Analysis of Whole-Genomes (PCAWG) project^34,35^, combined with almost all other cancer genomes and exomes for which suitable mutational catalogues are publicly available, https://www.synapse.org/#!Synapse:syn11801788, presents a notable opportunity to establish the repertoire of mutational signatures and to determine their activities across the range of cancer types.

## RESULTS

### Cancer genomes and somatic mutations

Somatic mutational catalogues from 23,829 samples of most cancer types, including the 2,780 highly curated PCAWG whole-genomes^34,35^, 1,865 additional whole-genomes and 19,184 exomes were studied. From these, 79,793,266 somatic single base substitutions, 814,191 doublet base substitutions and 4,122,233 small insertions and deletions (indels) were analysed for mutational signatures, ~10–fold more mutations than any previous study (https://www.synapse.org/#!Synapse:syn11801889)^4,36^.

To enable mutational signature analysis classifications were developed for each type of mutation. For single base substitutions, the primary classification comprised 96 classes constituted by the six base substitutions C>A, C>G, C>T, T>A, T>C, T>G (in which the mutated base is represented by the pyrimidine of the Watson-Crick base pair) plus the flanking 5’ and 3’ bases (https://cancer.sanger.ac.uk/cosmic/signatures/SBS). In some analyses, two flanking bases 5’ and 3’ to the mutated base were considered (generating 1,536 classes) or mutations within transcribed genome regions were selected and classified according to whether the pyrimidine of the mutated base pair fell on the transcribed or untranscribed strand (192 classes). A classification was also derived for doublet base substitutions (78 classes, https://cancer.sanger.ac.uk/cosmic/signatures/DBS). Indels were classified as deletions or insertions and, when of a single base, as C or T and according to the length of the mononucleotide repeat tract in which they occurred. Longer indels were classified as occurring at repeats or with overlapping microhomology at deletion boundaries, and according to the size of indel, repeat, and microhomology (83 classes, https://cancer.sanger.ac.uk/cosmic/signatures/ID).

### Mutational signature analysis

The mutational catalogues from the 2,780 PCAWG whole-genome, 1,865 additional whole-genome, and 19,184 exome sequences of cancer were analysed separately (https://doi.org/10.7303/syn11801889)^34,35^. For each of these catalogue sets, signature extraction was conducted using methods based on nonnegative matrix factorisation (NMF)^3,6^ on each cancer type individually and also on all cancer types together. Analyses were carried out separately for single base substitutions (SBS signatures), doublet base substitutions (DBS signatures) and indels (ID signatures) and also for the three mutation types together (1697 mutation classes if the 1536 classes of SBS in pentanucleotide context was employed) generating composite signatures.

Mutational signatures were extracted using two independently developed NMF-based methods: *(i)* SigProfiler, a further elaborated version of the framework used to generate the signatures shown in the previous version of the COSMIC compendium of mutational signatures (COSMICv2)^3,18,36–38^, and *(ii)* SignatureAnalyzer, based on a Bayesian variant of NMF used in several previous publications^6,15,39,40^. NMF determines both the signature profiles and the contributions of each signature to each cancer genome as part of its factorization of the input matrix of mutation spectra. However, given a substantial number of signatures and/or heterogeneous mutation burdens across samples, it is possible to reconstruct the mutations observed in a particular sample in multiple ways, often with very small and/or biologically implausible contributions from many signatures. Therefore, each method developed a separate procedure to estimate the contributions of signatures to each sample (Methods).

We tested SignatureAnalyzer and SigProfiler on 11 sets of synthetic data, encompassing a total of 64,400 synthetic samples, in which known signature profiles were used to generate catalogues of synthetic mutational spectra. Both approaches performed well in re-extracting the known signatures in realistically complex data. The tests highlighted the importance of, and challenges in, selecting the number of signatures, because extracted signatures discordant from the known input usually arose from difficulty in selecting the correct number of signatures. Thus, these tests confirmed that use of NMF-based approaches to extract signatures is not a purely algorithmic process. Instead, signature extraction requires human judgement that considers all of the available data, including evidence from experimental delineation of mutational signatures and the literature on DNA damage and repair, and prior evidence of biological plausibility. In addition, signature extraction requires human-guided sensitivity analysis to confirm that extractions from different groupings of tumours yield essentially the same signatures. These types of evidence and techniques were used in the determination of the signature profiles reported here. The findings we report from tests on synthetic data are consistent with results regarding NMF, and the related areas of probabilistic topic modelling and latent Dirichlet allocation, in multiple problem domains^41–43^. It is widely understood that the choice of the number of latent variables (for our purposes, the number of mutational signatures) is rarely amenable to complete automation. In further simulations, we also found that mutation catalogues from whole genomes allowed substantially better signature extraction than the much smaller catalogues from whole exomes and that signature extraction on whole genome data from half as many tumours would have supported inferior signature extraction. See Methods for further details; all results are at https://doi.org/10.7303/syn18497223 and a summary can be found at https://doi.org/10.7303/syn18511087.1.

The results of SigProfiler and SignatureAnalyzer exhibited many similarities, and we assigned the same identifiers to similar signatures extracted by the two methods https://www.synapse.org/#!Synapse:syn12016215. However, there were also noteworthy differences. The number of SBS signatures found in low mutation burden tumours in the PCAWG set (94.4% of cases that harbour 47% of mutations) was similar: 31 by SigProfiler and 35 by SignatureAnalyzer. The number of additional SBS signatures extracted from hyper-mutated PCAWG samples (5.6% of cases and 53% of mutations), however, was different: 13 by SigProfiler and 25 by SignatureAnalyzer. There were also differences in SBS signature profiles, including among signatures found in low mutation burden cases. The latter primarily involved “flat”, relatively featureless signatures, which are mathematically challenging to deconvolute. Finally, there were differences in signature attributions to individual samples. In general, SignatureAnalyzer used more signatures to reconstruct the mutational profiles (Extended Data Figure 1, https://www.synapse.org/#!Synapse:syn12169204, https://www.synapse.org/#!Synapse:syn12177011) and the attribution to flat signatures was different, with SigProfiler assigning mutations to SBS5 and SBS40 and SignatureAnalyzer using combinations of multiple signatures (Extended Data Figure 2ab, https://www.synapse.org/#!Synapse:syn12169204). The DBS and ID signatures were generally similar between the two methods (Extended Data Figure 2cd). These comparisons provide a useful perspective on both the consistency and variability of signature extraction and attribution depending on the methodology used.

The final sets of reference mutational signatures were determined from the PCAWG analysis supplemented by additional signatures from the other datasets. SBS signatures using the 96 mutation classification were supported by the outcomes of analyses using the 192 and 1536 mutation classifications, the existence of individual cancer samples dominated by a particular signature, and, where available, prior experimental evidence for certain mutational signatures (Methods, https://doi.org/10.7303/syn12025148, https://doi.org/10.7303/syn12009645, COSMIC at https://cancer.sanger.ac.uk/cosmic/signatures). Each signature was allocated a number consistent with, and extending, the COSMICv2 annotation^37^. Some previous signatures split into multiple constituent signatures and these were numbered as before but with additional letter suffixes (e.g., single SBS17 split into signatures SBS17a and SBS17b). DNA sequencing and analysis artefacts also generate mutational signatures, and we indicate which signatures are possible artefacts (https://www.synapse.org/#!Synapse:syn12009767) but do not present them below. However, future studies employing this signature set as a reference may consider utilizing artefact signatures for data quality control. The results of both SignatureAnalyzer and SigProfiler were used throughout the research reported here. However, for brevity and for continuity with the signature set previously displayed in COSMIC^37^, which has been widely used as a reference, SigProfiler results are outlined below and SignatureAnalyzer results are provided at (Extended Data Figures 3,4, https://www.synapse.org/#!Synapse:syn11738307).

### Single base substitution (SBS) mutational signatures

There were substantial differences in numbers of SBSs between samples (ranging from hundreds to millions) and between cancer types, as previously observed^44^ (Figure 1). In total, 67 SBS mutational signatures were extracted, of which 49 were considered to be likely of biological origin (Figure 2, Methods, https://cancer.sanger.ac.uk/cosmic/signatures/SBS/). Except for SBS25, all mutational signatures reported in COSMICv2 (i.e., https://cancer.sanger.ac.uk/cosmic/signatures_v2) ^4,23,37^ were confirmed in the new set of analyses (median cosine similarity between the newly derived signatures and those on COSMICv2: 0.95, excluding "split" signatures which are discussed below; range 0.74 to 0.9996 https://www.synapse.org/#!Synapse:syn12016215). SBS14, SBS16, and SBS20 changed the most; for explanation, see https://cancer.sanger.ac.uk/cosmic/signatures/SBS/. SBS25 was previously found only in cell lines derived from Hodgkin lymphomas, at least one of which had been previously treated with chemotherapy, and, to our knowledge, no data from primary cancers of this type are currently available. The newly derived signatures showed much improved separation from each other and hence more distinct signature profiles, presumably due to the substantially increased statistical power of this analysis (online Methods section *Better separation compared to COSMICv2 signatures*).

**Figure 1.**
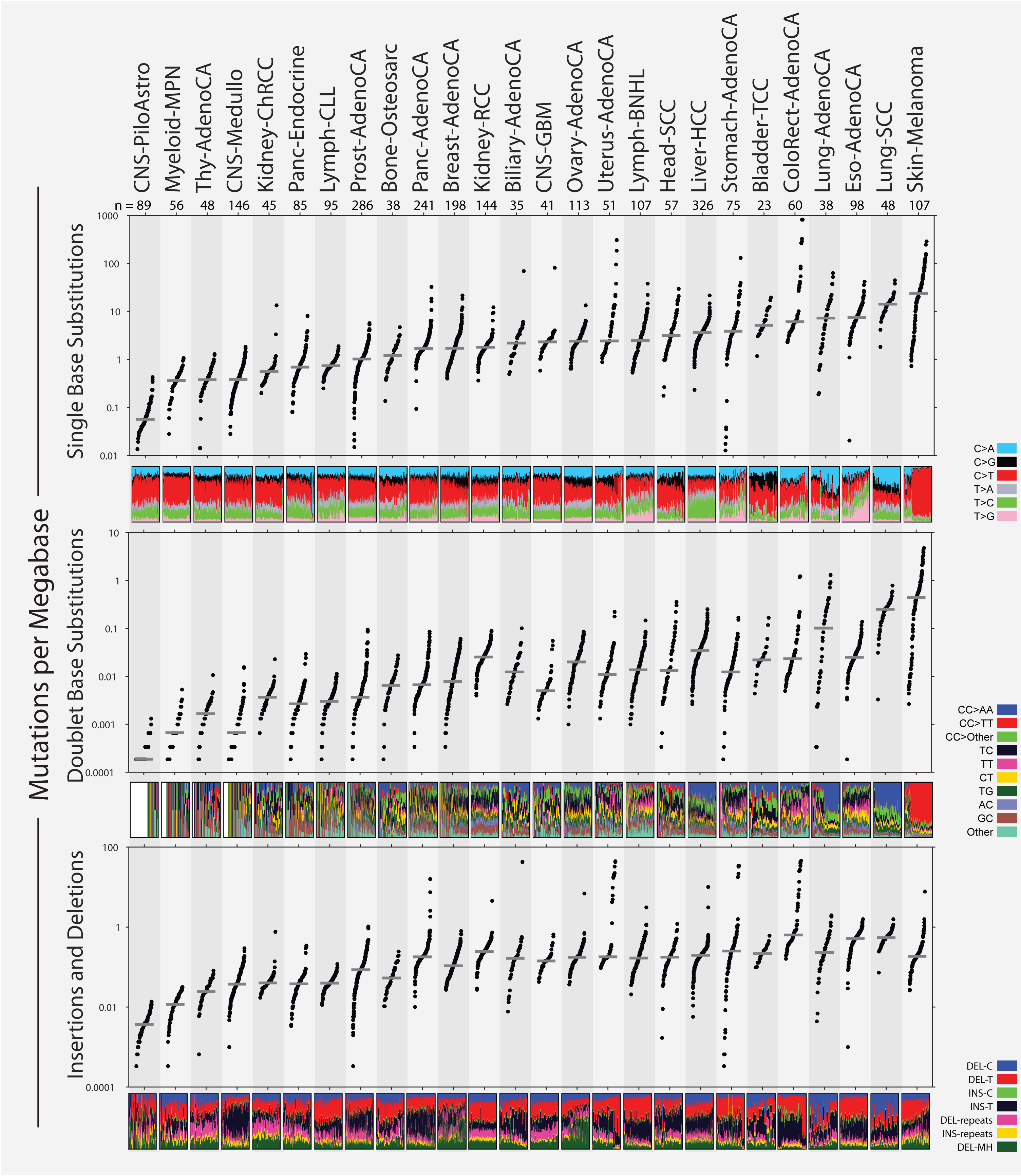
Mutation burdens of single base substitutions, doublet base substitutions and small insertions and deletions for the 2,780 PCAWG tumours. Each sample is displayed according to its tumour type. Tumour types are ordered according to the median number of single base substitutions. The numbers of cases of each tumour type are shown. The proportions of each mutation subclass in each sample are shown as coloured bar charts.

**Figure 2.**
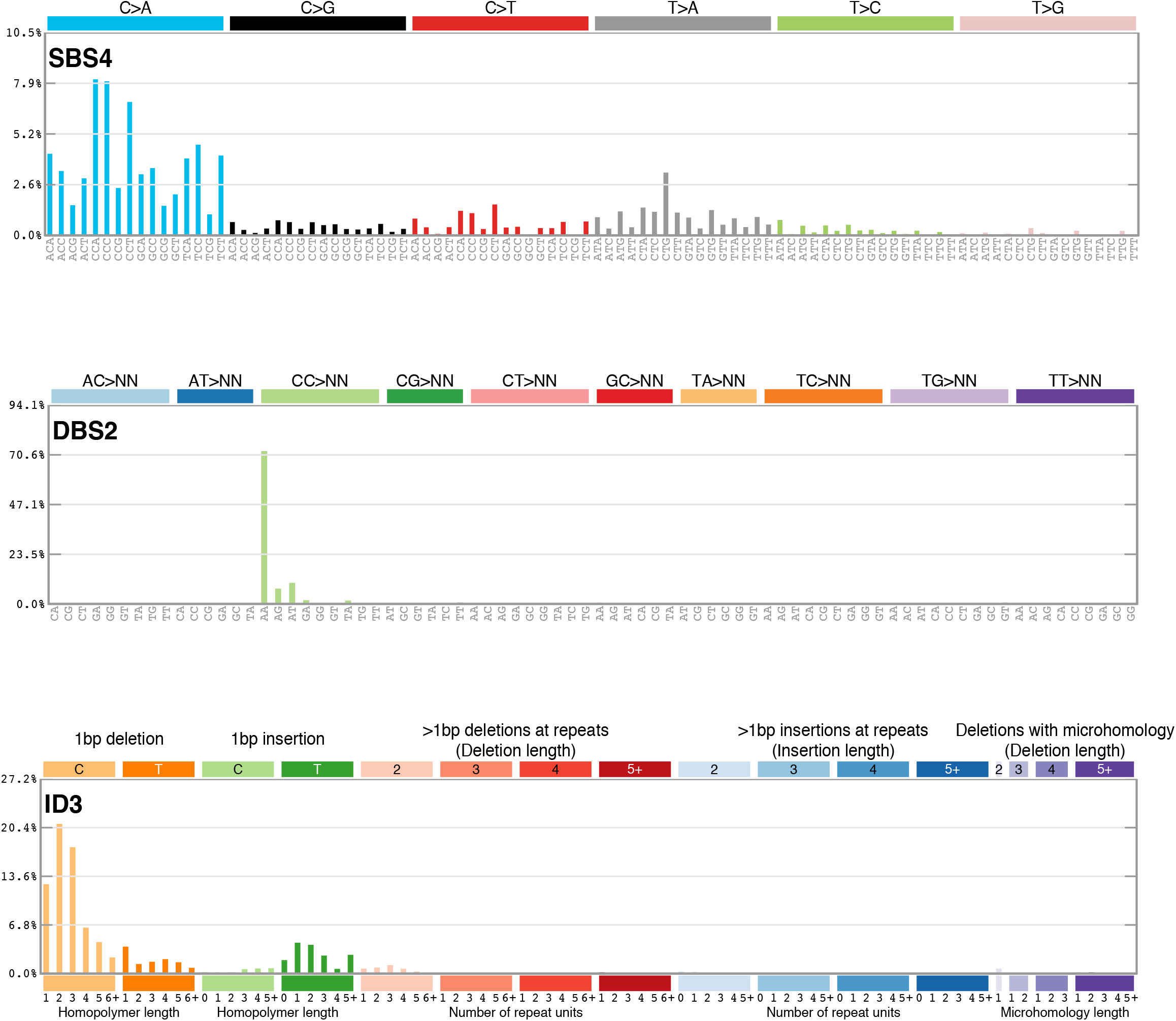
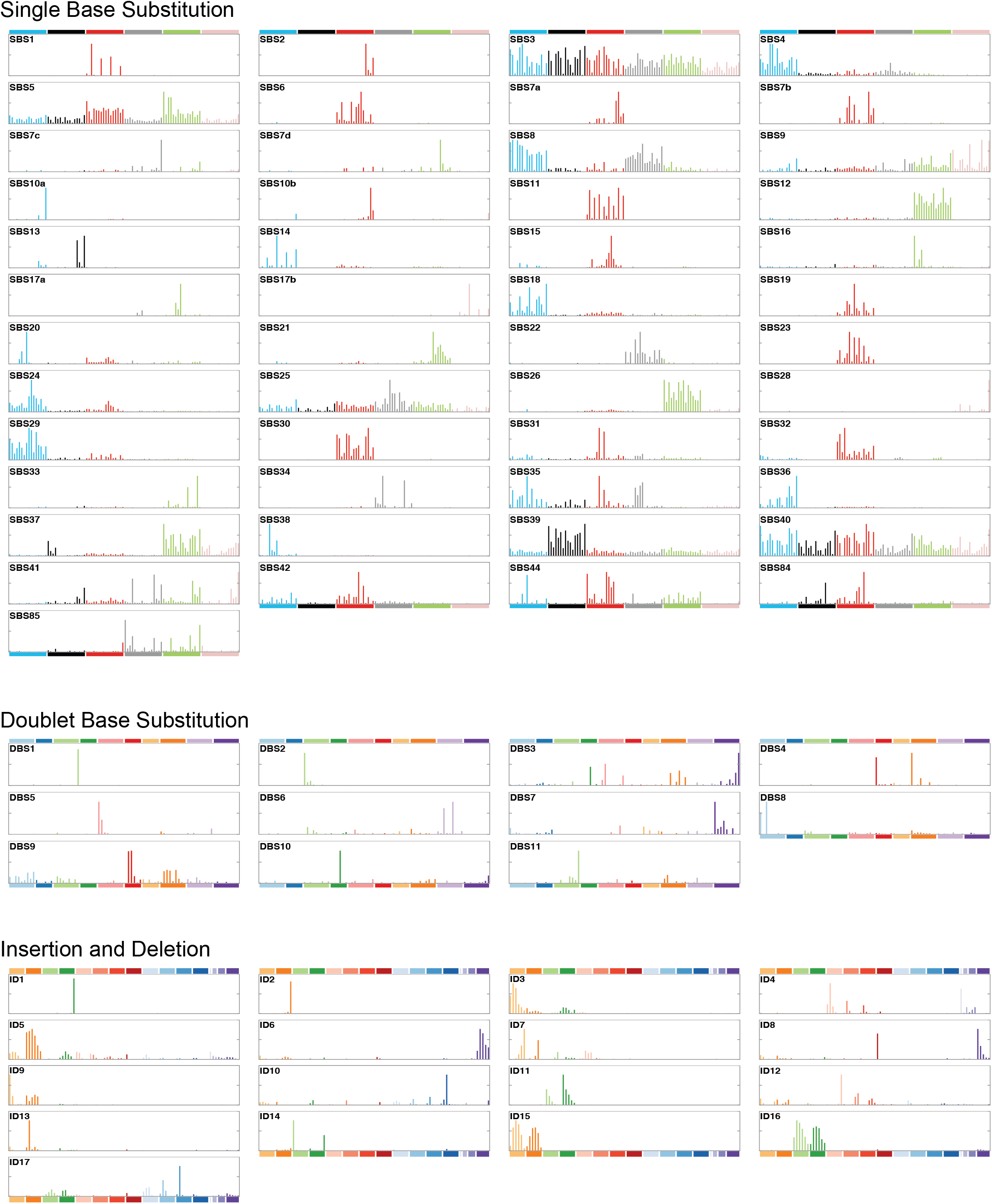
Profiles of single base substitution, doublet base substitution and small insertion and deletion mutational signatures. The subclassifications of each mutation type (single base substitutions, 96 subtypes; doublet base substitutions, 78 subtypes; indels, 83 subtypes) are described in the main text. Magnified versions of signatures SBS4, DBS2 and ID3 (which are all associated with tobacco smoking) are shown to illustrate the positions of each mutation subtype on each plot.

Thirteen new likely real SBS signatures compared to the set previously described in COSMICv2^37^ were extracted (excluding those that are the consequence of signature splitting). Some were in cancers with a previously unanalysed exogenous exposure (SBS42), some were in chemotherapy treated samples which have often been excluded from previous studies (SBS31, SBS32, SBS35) and some were rare and hence absent by chance from previous analyses (SBS36, SBS44). Others were more common, but contributed relatively few mutations to individual cancer genomes, or were similar to previously discovered signatures and thus not isolated from datasets based predominantly on cancer exome sequences (e.g., SBS38, SBS39, SBS40). Notably, SBS40 was extracted from kidney cancer in which it appears to be required for optimal reconstruction of mutational catalogues. It is a relatively featureless (“flat”) signature, with similarity to SBS5 and other flat signatures, and this may account for it only clearly emerging now with the availability of whole cancer genomes. SBS40 may contribute to other cancer types but its similarity to SBS5 renders this uncertain and larger datasets will be required to clarify the extent of its activity. For some new signatures there were plausible underlying aetiologies (Figure 3, Extended Data Figures 4,5): SBS31 and SBS35, prior platinum compound chemotherapy^45^; SBS32, prior azathioprine therapy; SBS36, inactivating germline or somatic mutations in *MUTYH* which encodes a component of the base excision repair machinery^46,47^; SBS38, additional effects of ultraviolet light (UV) exposure; SBS42, occupational exposure to haloalkanes^27^; SBS44, defective DNA mismatch repair due to MLH1 inactivation^48^. SBS33, SBS34, SBS37, SBS39, SBS40, and SBS41 are of unknown cause.

**Figure 3.**
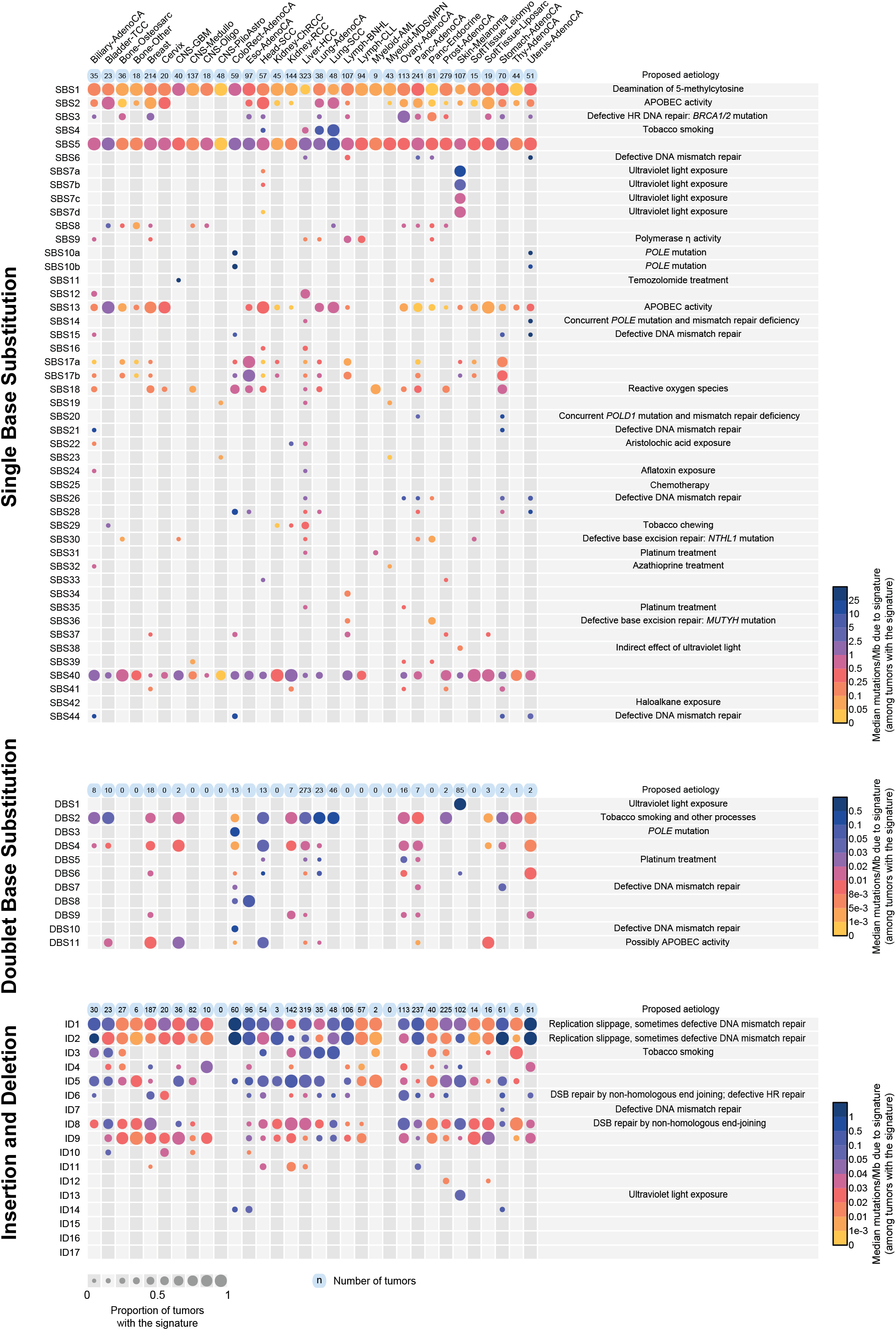
The number of mutations contributed by each mutational signature to the 2,780 PCAWG tumours. The numbers of mutations attributed are shown by cancer type. The size of each dot represents the proportion of samples of each tumour type that show the mutational signature. The colour of each dot represents the median mutation burden of the signature in samples which show the signature. Contributions are shown for single base substitution, doublet base substitution and indel mutational signatures separately. Contributions of composite signatures to the PCAWG cancers and single base substitution signatures to the complete set of cancer samples analysed are shown in Supplementary information.

Three previously characterised base substitution signatures (SBS7, SBS10, SBS17) split into multiple constituent signatures (Figure 2). We previously regarded SBS7 as a single signature composed predominantly of C>T at C<u>C</u>N and T<u>C</u>N trinucleotides (the mutated base is underlined) together with many fewer T>N mutations. It was found in malignant melanomas and squamous skin carcinomas and is likely due to UV induced pyrimidine dimer formation followed by translesion DNA synthesis by error-prone polymerases which predominantly insert adenine opposite damaged bases. With the larger dataset now available, SBS7 has decomposed into four constituent signatures: SBS7a consisting mainly of C>T at T<u>C</u>N; SBS7b consisting of C>T mainly at C<u>C</u>N and to a lesser extent at T<u>C</u>N; SBS7c and SBS7d, which constituted relatively minor components of the previous SBS7 and consist predominantly of T>A at N<u>T</u>T and T>C at N<u>T</u>T respectively^49^. Splitting of a mutational signature likely reflects the existence of multiple distinct mutational processes, initiated by the same exposure, which have closely, but not perfectly, correlated activities. For example, the constituent signatures of SBS7 are probably all initiated by UV-induced DNA damage. SBS7a and SBS7b may reflect different dipyrimidine photoproducts whereas SBS7c and SBS7d may be due to low frequencies of misincorporation by translesion polymerases of T and G opposite thymines in pyrimidine dimers rather than the more frequent and non-mutagenic A. Splitting of SBS10 and SBS17 is described at https://cancer.sanger.ac.uk/cosmic/signatures/SBS/.

Several base substitution signatures showed transcriptional strand bias (https://www.synapse.org/#!Synapse:syn12009767). Transcriptional strand bias is often attributable to transcription coupled nucleotide excision repair (TC-NER) acting on DNA damaged by exogenous exposures which cause covalently bound bulky adducts or crosslinking to other bases and consequent distortion of the helical structure. This results in stalling of RNA polymerase and hence recruitment of the TC-NER machinery. An excess of DNA damage on untranscribed compared to transcribed strands of genes may also contribute to transcriptional strand bias^50^. Both mechanisms, however, result in more mutations of a damaged base on the untranscribed compared to the transcribed strands of genes. Assuming that either or both are responsible for the observed transcriptional strand biases (which may not always be the case), DNA damage to cytosine (SBS7a, SBS7b), guanine (SBS4, SBS8, SBS19, SBS23, SBS24, SBS31, SBS32, SBS35, SBS42), thymine (SBS7c, SBS7d, SBS21, SBS26, SBS33) and adenine (SBS5, SBS12, SBS16, SBS22, SBS25) may underlie these mutational signatures (see https://cancer.sanger.ac.uk/cosmic/signatures/SBS/ for plots of strand bias). Although the likely underlying DNA damaging agents are known for SBS4 (tobacco mutagens), SBS7a, SBS7b, SBS7c, SBS7d (UV), SBS22 (aristolochic acid), SBS24 (aflatoxin), SBS25 (prior chemotherapy), SBS31 and SBS35 (platinum compounds), SBS32 (azathioprine), and SBS42 (haloalkanes), the causes of the remainder are unknown. Indeed, some signatures showing transcriptional strand bias are associated with defective DNA mismatch repair (SBS21 and SBS26) and it is conceivable that, for these, exogenous DNA damage is not involved. The extent of transcriptional strand bias appears to differ in different sectors of the genome. For example, consideration of the whole transcribed genome showed absent or minimal transcriptional strand bias in the APOBEC related SBS2 and SBS13 and in the defective polymerase epsilon proof-reading related SBS10a. However, consideration of exons alone showed clear evidence of transcriptional strand bias in these signatures (https://cancer.sanger.ac.uk/cosmic/signatures/SBS/). The mechanism(s) underlying this amplification of transcriptional strand bias in exons is unknown and appears to be signature specific, since there is minimal difference in the extent of transcriptional strand bias between exons and other transcribed regions for other signatures (for example, SBS4 and SBS22).

Employing the single base substitution classification of 1536 mutation types, which uses the pentanucleotide sequence context two bases 5’ and two bases 3’ to each mutated base, yielded a set of signatures largely consistent with that based on substitutions in trinucleotide context alone. Notably, however, the pentanucleotide context enabled the extraction of two forms of both SBS2 and SBS13, one with mainly a pyrimidine (C or T) and the other with a purine (A or G) at the −2 base (the second base 5’ to the mutated cytosine). These may represent the activities of the cytidine deaminases APOBEC3A and APOBEC3B, respectively^51^. If so, APOBEC3A accounts for many more mutations than APOBEC3B in cancers with high APOBEC activity. Several other signatures showed non-random sequence contexts at +2 and −2 positions. In particular, the −2 bases in SBS17a and SBS17b and the −2 and +2 bases in SBS9 were predominantly A and T. In general, however, sequence context effects were much stronger for bases immediately 5’ and 3’ to the mutated bases.

SBS signatures showed substantial variation in the numbers of cancer types and cancer samples in which they were found, ranging from SBS1 and SBS5 which were present in almost every cancer type and almost every cancer sample, to SBS23 which was only observed in a small subset of liver cancers (Figure 3). The numbers of mutations per cancer sample attributed to each signature also varied greatly, from a few tens of mutations for SBS1 to millions of mutations for SBS10b. Almost all individual cancer samples exhibited multiple signatures, with a mode of three signatures per sample in the PCAWG set (https://www.synapse.org/#!Synapse:syn12169204). The assigned signatures reconstruct well the mutational spectra of the cancer samples (in PCAWG samples, median cosine similarity 0.97; 96.3% of samples with cosine similarity >0.90) (illustrative examples are shown in Figure 4).

**Figure 4.**
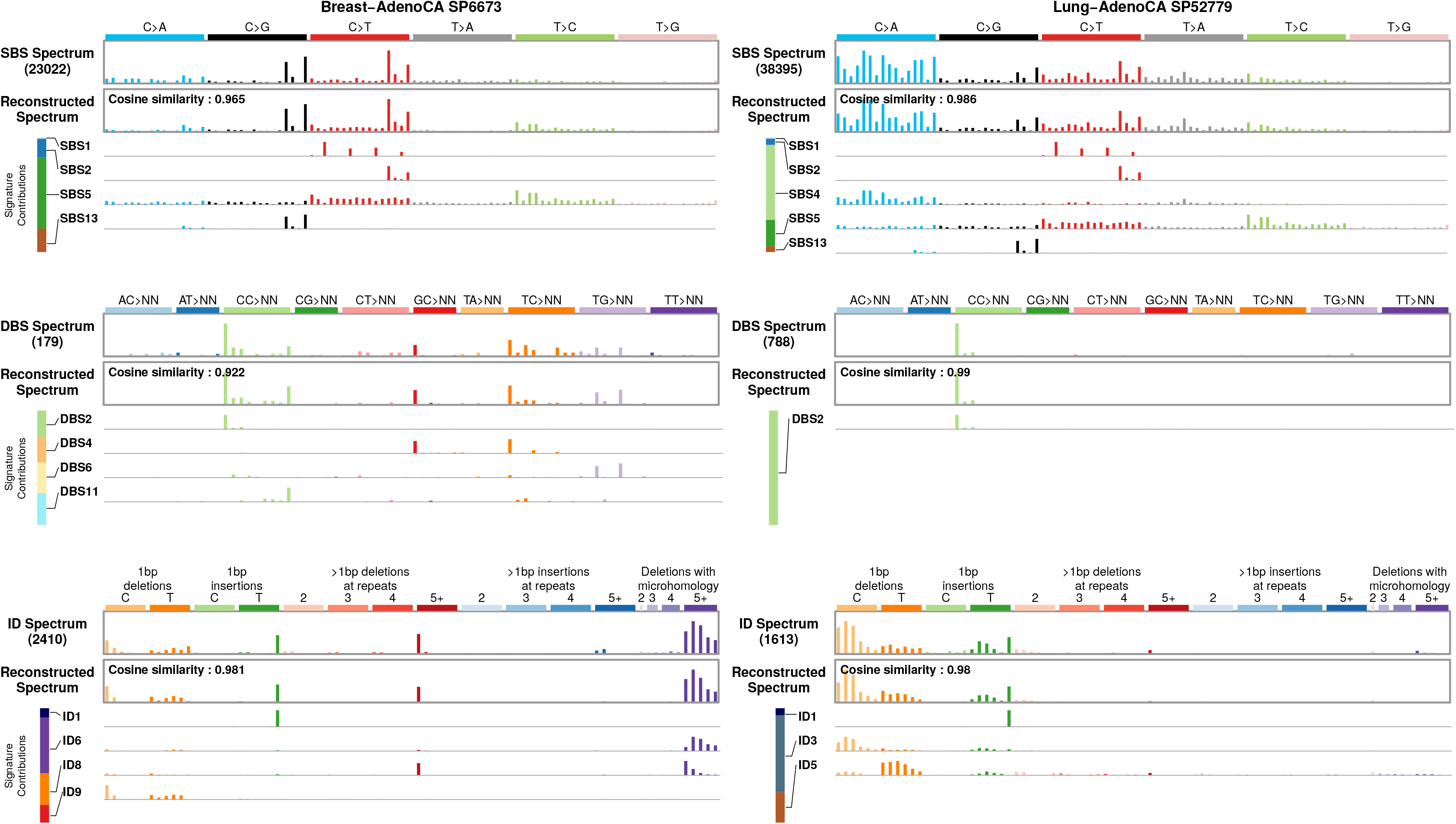
Illustrative examples of mutational spectra of individual cancer samples. A breast cancer, a lung cancer, and a malignant melanoma and their contributory single base substitution, doublet base substitution, and small insertion and deletion mutational signatures.

### Clustered single base substitution mutational signatures

Some mutational processes generate mutations that cluster in small regions of the genome. The relatively limited number of mutations generated by such processes, compared to those acting genome-wide, may result in failure to detect their signatures by standard methods. To obviate this problem, we first identified clustered mutations in each genome and analysed these separately (Methods). Four main signatures associated with clustered mutations were identified (Figure 2) and were consistent with previous reports^15,16,32^. Two found in multiple cancer types were similar to single base substitution SBS2 and SBS13, which have been attributed to APOBEC enzyme activity (mostly APOBEC3B) and represent foci of *kataegis*^17,32,52^. Two additional clustered mutational signatures, one characterised by C>T and C>G mutations at (A|G)<u>C</u>(C|T) trinucleotides^53^ and the other T>A and T>C mutations at (A|T)<u>T</u>(A|T) were found in lymphoid neoplasms and likely represent direct and indirect consequences of activation induced cytidine deaminase (AID) mutagenesis and translesion DNA synthesis by error-prone polymerases (SBS84 and SBS85 respectively)^15^. The possibility that further processes may generate clustered mutations is not excluded.

### Doublet base substitution (DBS) mutational signatures

Tandem doublet, triplet, quadruplet, quintuplet, and sextuplet base substitutions (https://www.synapse.org/#!Synapse:syn11801938, https://www.synapse.org/#!Synapse:syn11726620) at immediately adjacent bases were observed at ~1% the prevalence of single base substitutions. In most cancer genomes, the observed number of DBSs was considerably higher than expected from random adjacency of SBSs (https://www.synapse.org/#!Synapse:syn12177057) indicating the existence of commonly occurring, single mutagenic events that cause substitutions at neighbouring bases. There was substantial variation in the number of DBSs, ranging from zero to 20,818 in a sample. Across cancer types, the numbers of DBSs were generally proportional to the numbers of SBSs in that cancer type (Figure 1). However, colorectal adenocarcinomas had significantly fewer DBSs than expected, and lung cancers and melanomas had more (Extended Data Table 1). The large dataset analysed here allowed, for the first time, systematic analysis of DBS and indel signatures (described below). Eleven DBS signatures were extracted (Figure 2). Of these, to our knowledge, only two have been previously reported^33^ evidencing further the value of the large numbers of mutations from whole genome data.

DBS1 was characterised almost exclusively by CC>TT mutations (Figure 2), contributed 100s-10,000s of mutations in malignant melanomas (Figure 3) with SBS7a and SBS7b. DBS1 exhibited transcriptional strand bias consistent with damage to cytosines (https://www.synapse.org/#!Synapse:syn12177063). CC>TT mutations associated with UV induced DNA damage are well established in the literature, were previously reported in melanomas, and are thought to be due to generation of pyrimidine dimers and subsequent error-prone translesion DNA synthesis by polymerases that introduce adenines opposite the damaged bases^33,54^.

Reanalysis after exclusion of malignant melanomas and other cancers with evidence of UV exposure still yielded a signature (termed DBS11) characterised predominantly by CC>TT mutations and smaller numbers of other doublet base substitutions at CC and TC which contributed 10s of mutations to many samples of multiple cancer types (Figures 2 and 3). DBS11 was associated with SBS2 which is due to APOBEC activity. Thus, APOBEC activity may also generate DBS11, although the mechanism by which it induces doublet base substitutions is not well understood.

DBS2 was composed predominantly of CC>AA mutations, with smaller numbers of CC>AG and CC>AT mutations, and contributed 100s-1000s of mutations in lung adenocarcinoma, lung squamous and head and neck squamous carcinomas, which are often caused by tobacco smoking, which has been reported previously (Figures 2 and 3)^33^. DBS2 showed transcriptional strand bias indicative of guanine damage (https://www.synapse.org/#!Synapse:syn12177064) and was associated with SBS4 which is caused by tobacco smoke exposure. It is likely, therefore, that DBS2 can be a consequence of DNA damage by tobacco smoke mutagens.

Analysis of each cancer type separately, however, revealed a signature very similar to DBS2 contributing 100s of mutations to liver cancers and 10s of mutations to cancers of other types without evidence of tobacco smoke exposure. A pattern closely resembling DBS2 and characterised predominantly by CC>AA mutations, together with smaller contributions of CC>AG and CC>AT, dominates DBSs in normal mouse cells and is particularly frequent in the liver^55^. The nature of the mutational processes underlying these doublet signatures in smoking-unrelated human cancers and in normal mice is unknown. However, acetaldehyde exposure in experimental systems generates a mutational signature characterised primarily by CC>AA and lower burdens of CC>AG and CC>AT mutations together with C>A single base substitutions^56^. Acetaldehyde is an oxidation product of alcohol and a constituent of cigarette smoke. The role of acetaldehyde, and perhaps other aldehydes, in generating DBS2, whether associated with tobacco smoking, alcohol consumption or in non-exposed cells, merits further investigation^57^.

DBS3, DBS7, DBS8 and DBS10 showed 100s-1000s of mutations in rare colorectal, stomach and oesophageal cancers some of which showed evidence of defective DNA mismatch repair (DBS7, DBS10) or polymerase epsilon exonuclease domain mutations (DBS3) generating hypermutator phenotypes (Figures 2, 3). DBS5 was found in cancers previously exposed to platinum chemotherapy and is associated with SBS31 and SBS35. The remaining DBS signatures are of uncertain cause.

### Small insertion and deletion (ID) mutational signatures

Indels were usually present at ~10% the frequency of base substitutions (Figure 1). There was substantial variation between cancer genomes in numbers of indels, even when cancers with evidence of defective DNA mismatch repair were excluded. Overall, the numbers of deletions and insertions were similar, but there was variation between cancer types with some showing more deletions and others more insertions of various subtypes (Figure 1). Seventeen indel mutational signatures were extracted (Figure 2).

Indel signature 1 (ID1) was composed predominantly of insertions of thymine and ID2 of deletions of thymine, both at long (≥5) thymine mononucleotide repeats (Figure 2). 10s to 100s of mutations of both signatures were found in the large majority of most cancer types but were particularly common in colorectal, stomach, endometrial and oesophageal cancers and in diffuse large B cell lymphoma (Figure 3). Most of these cancers are likely to be DNA mismatch repair proficient on the basis of the relatively limited numbers of indels and absence of the SBS signatures (SBS6, SBS14, SBS15, SBS20, SBS21, SBS26, and SBS44) associated with DNA mismatch repair deficiency. Together, ID1 and ID2 accounted for 97% and 45% of indels in hypermutated and non-hypermutated cancer genomes, respectively (Extended Data Table 2), and both signatures have also been found in non-neoplastic cells^58^. They are likely due to the intrinsic tendency to slippage during DNA replication of long mononucleotide tracts. However, the mechanistic basis for separation into two signatures, one presumably due to slippage of the nascent strand (ID1) and the other the template strand (ID2) is unclear. Similarly, the substantial differences in their mutation frequencies between cancer types are not well understood.

ID3 was characterised predominantly by deletions of cytosine at short (≤5bp long) mononucleotide cytosine repeats and exhibited 100s of mutations in tobacco smoking associated cancers of the lung and head and neck (Figures 2 and 3). There was transcriptional strand bias of mutations, with more guanine deletions than cytosine deletions on the untranscribed strands of genes, compatible with TC-NER of adducted guanine (https://www.synapse.org/#!Synapse:syn12177065, https://www.synapse.org/#!Synapse:syn12177066). The numbers of ID3 mutations in cancer samples positively correlated with the numbers of SBS4 and DBS2 mutations, both of which have been associated with tobacco smoking (Extended Data Figure 6). It is therefore likely that DNA damage by components of tobacco smoke underlie ID3 but the mechanism(s) by which indels are generated is unclear.

ID13 was characterised predominantly by deletions of thymine at thymine-thymine dinucleotides and exhibited large numbers of mutations in malignant melanomas of the skin (Figures 2 and 3). The numbers of ID13 mutations correlated with the numbers of SBS7a, SBS7b and DBS1 mutations, which have been attributed to DNA damage induced by UV (Extended Data Figure 6). It is, however, notable that a similar mutation of the other pyrimidine, i.e., deletion of cytosine at cytosine-cytosine dinucleotides, does not feature strongly in ID13, perhaps reflecting the predominance of thymine compared to cytosine dimers induced by UV^59^. The mechanism(s) underlying thymine deletion is unclear.

ID6 and ID8 were both characterised predominantly by deletions ≥5bp (Figure 2). ID6 exhibited overlapping microhomology at deletion boundaries with a mode of 2bp and often longer stretches. This signature was correlated with SBS3 which has been attributed to defective homologous recombination based repair (Extended Data Figure 6). By contrast, ID8 deletions showed shorter or no microhomology at deletion boundaries, with a mode of 1bp, and did not strongly correlate with SBS3 mutations (Figures 2 and 3). These patterns of deletion may be characteristic of DNA double strand break repair by non-homologous recombination based end-joining mechanisms, and if so, suggest that at least two distinct forms of end-joining mechanism are operative in human cancer^60^.

A small fraction of cancers exhibited very large numbers of ID1 and ID2 mutations (>10,000) (Figure 3, https://cancer.sanger.ac.uk/cosmic/signatures/ID). These were usually accompanied by SBS6, SBS14, SBS15, SBS20, SBS21, SBS26 and/or SBS44 which are associated with DNA mismatch repair deficiency, sometimes combined with POLE or POLD1 proofreading deficiency (SBS14, SBS20)^40^. Occasional cases with these signatures additionally showed large numbers of ID7 indels (https://www.synapse.org/#!Synapse:syn11738668). In addition, rare samples showed large numbers of either ID4, ID11, ID14, ID15, ID16 or ID17 mutations but did not show ID1 and ID2 mutations or the single base substitution signatures usually associated with DNA mismatch repair deficiency. The mechanisms underlying these signatures are unknown.

### Composite mutational signatures

In the analyses described above mutational signatures were extracted for each mutation type separately. However, mutational processes in nature generate composite signatures that may include SBSs, DBSs, IDs, genome rearrangements and chromosome number changes. We therefore also extracted signatures using combined catalogues of SBSs, DBSs, and IDs (257 mutation subclasses or 1697 if the 1536 classification of single base substitutions was used). Fifty-two composite signatures were extracted.

A composite signature with components similar to SBS4, DBS2 (characterised predominantly by CC>AA mutations) and ID3 (characterised predominantly by deletion of cytosine at short runs of cytosines) was found mainly in lung cancers, suggesting that it is the consequence of tobacco smoke exposure (Extended Data Figure 7). Similarly, composite signatures with components similar to SBS7a, SBS7b, DBS1 (characterised predominantly by CC>TT mutations) and ID13 (characterised predominantly by deletion of thymine at thymine– thymine dinucleotides) were found in skin cancers and are thus likely due to UV induced DNA damage (Extended Data Figure 7). A further composite signature in breast and ovarian cancers included features of SBS3 and ID6 combined with ID8 (deletions >5bp with varying degrees of overlapping microhomology) and is likely associated with defective homologous recombination based repair (Extended Data Figure 7). In these composite signatures attributions of the constituent SBS, DBS and ID signatures extracted independently in the main analyses were correlated with each other, adding support to the existence of the composite signatures (Extended Data Figure 6). Various forms of defective DNA mismatch repair were also associated with multiple SBS, DBS and ID signatures.

### Correlations with age

A positive correlation between age of cancer diagnosis and the number of mutations attributable to a signature suggests that the mutational process underlying the signature has been operative, at a more or less constant rate, throughout the cell lineage from fertilized egg to cancer cell, and thus in normal cells from which that cancer type develops^4,61^. Confirming previous reports, the numbers of SBS1 and SBS5 mutations correlated with age, exhibiting different rates in different tissue types (q-values in https://www.synapse.org/#!Synapse:syn12030687, https://www.synapse.org/#!Synapse:syn20317940 https://www.synapse.org/#!Synapse:syn12217988). In addition, SBS40 correlated with age in multiple cancer types. However, given the similarity in signature profile between SBS5 and SBS40 the possibility of misattribution between these signatures cannot currently be excluded. The numbers of DBSs and IDs were much lower than the numbers of SBSs and the numbers of samples in which DBS and ID signatures could be attributed were also lower. Nevertheless, DBS2 and DBS4 correlated with age and, consistent with the interpretation of activity in normal cells, the profiles of DBS2 and DBS4 together closely resemble the spectrum of DBS mutations found in normal mouse cells^55^. Neither DBS2 nor DBS4, however, was clearly correlated with an SBS or ID signature that correlates with age. ID1, ID2, ID5 and ID8 showed correlations with age in multiple tissues. ID1 and ID2 indels are likely due to slippage at poly T repeats during DNA replication and correlated with the number of SBS1 substitutions. SBS1 has previously been proposed to reflect the number of mitoses a cell has experienced and thus SBS1, ID1 and ID2 may all be generated during DNA replication at mitosis^4^. The number of ID5 mutations correlated with the number of SBS40 mutations and thus the mutational processes underlying these two age-correlated signatures may also harbour common components. ID8 is predominantly composed of deletions >5bp with no or 1bp of microhomology at their boundaries. These are likely due to DNA double strand breaks which have not been repaired by homologous recombination based mechanisms, but instead by a non-homologous-end joining mechanism. The features of ID8 resemble those of some ionising radiation associated mutations and this may, therefore, be an underlying aetiological factor^62^. Taken together, the results indicate that multiple mutational processes operate in normal cells.

## DISCUSSION

Cancers arise as a result of somatic mutations. Mutational signature analysis therefore provides important insights into cancer development through comprehensive characterisation of the underlying mutational processes. There are, however, important constraints, limitations and assumptions in the analytic frameworks we have used that should be recognised. Although designed to reflect the mutational consequences of recurrent mutational processes, mutational signatures extracted from sample sets in which multiple mutational processes are operative remain mathematical approximations, with profiles that can be influenced by the mathematical approach used and by additional factors, such as the other mutational processes present. For conceptual and practical simplicity, we have assumed that there is a single signature associated with each mutational process and have provided an average reference signature to represent it. However, we do not discount the possibility that further nuances and variations of signature profiles exist, for example between different tissues. Moreover, although the extent of separation between partially correlated signatures has been improved in this analysis, some signatures may still represent combinations of constituent signatures. Contributions from each signature to the burden of mutations in each sample have been estimated. However, with increasing numbers of signatures and multiple orders of magnitude differences in mutation burdens from certain signatures, prior knowledge can help to avoid biologically implausible results. Thus, further development of methods for deciphering mutational signatures and attribution of mutations is warranted and this needs to be supplemented by signatures derived from experimental systems in which the causes of the mutations are known. The numbers of DBSs, clustered substitutions, IDs and genome rearrangements (reported in ref. ^30^) are small compared to single base substitutions. Thus, larger datasets may be required to robustly characterise their mutational signatures. Nevertheless, the results outlined here indicate that signatures with many similarities and some differences can be found by different mathematical approaches, and that these are confirmed in many different ways, including experimentally elucidated signatures^22,31,45,48,49,61,63–69^ and the observation of tumours dominated by a single signature (https://www.synapse.org/#!Synapse:syn12016215).

Prior reports have provided only a relatively limited examination of doublet and indel mutational spectra and, to the best of our knowledge, no previous comprehensive analysis of doublet and indel mutational signatures has been performed. Here, we provide the first systematic analysis of these mutation types by considering 83 mutational subtypes for indels and 78 mutational subtypes for doublets. This analysis also includes almost all publicly available exome and whole-genome cancer sequences, amounting in aggregate to 23,829 cancers of most cancer types. Some rare or geographically restricted signatures may not have been captured and signatures of therapeutic mutagenic exposures have not been exhaustively explored. Nevertheless, it is likely that a substantial proportion of the naturally-occurring mutational signatures found in human cancer have now been described. This comprehensive repertoire provides a foundation for future research into *(i)* geographical and temporal differences in cancer incidence to elucidate underlying differences in aetiology, *(ii)* the mutational processes and signatures present in normal tissues and caused by non-neoplastic disease states, *(iii)* clinical and public health applications of signatures as indicators of sensitivity to therapeutics and past exposure to mutagens, and *(iv)* mechanistic understanding of the mutational processes underlying carcinogenesis.

## ACKNOWLEDGEMENTS

The results here are partly based on data generated by the TCGA Research Network (http://cancergenome.nih.gov/) and the ICGC/TCGA Pan-cancer Analysis of Whole Genomes Network. This work was supported by Wellcome grant reference 206194 (M.R.S.), Singapore National Medical Research Council grants NMRC/CIRG/1422/2015 and MOH-000032/MOH-CIRG18may-0004 and the Singapore Ministry of Health via the Duke-NUS Signature Research Programmes (M.N.H., A.W.T.N., Y.W., A.B., S.G.R.), US National Institute of Health Intramural Research Program Project Z1AES103266 (D.A.G.), the European Research Council Consolidator Grant 682398 (N.L.-B.), US National Cancer Institute U24CA143843 (D.A.W.), and Cancer Research UK Grand Challenge Award C98/A24032 (E.N.B., S.M.A.I., L.B.A., M.R.S.). G.G and J.K were partially supported by the National Cancer Institute grants U24CA210999 and U24CA143845. G.G. was partially supported by the Paul C. Zamecnick, MD, Chair in Oncology at the Massachusetts General Hospital Cancer Center. N.J.H and G.G were partially supported by G.G’s start up funds at MGH.

## Online Methods

### Principles and strategy of mutational signature analysis adopted in this report

#### Conceptual principles

- Multiple mutational processes generate the somatic mutations present in each individual human cancer.
- Each mutational process generates a particular pattern of somatic mutations known as a mutational signature.
- Each mutational process may incorporate a component of DNA damage/modification, DNA repair and DNA replication, each of which may be part of normal or abnormal cell biology. Differences in any of the three components may result in a different mutational signature, thus, by definition, constituting a distinct mutational process.
- Multiple mutational processes operating continuously or intermittently during the cell lineage from the fertilised egg to the cancer cell may contribute to the aggregate set of mutations found in the cancer cell. Thus, the catalogue of somatic mutations from a single cancer sample often includes mutations of many different mutational signatures.

#### Aims of the study

- To decipher the mutational signatures present in essentially the full set of whole genome and exome sequenced human cancers from which data is currently available and subsequently to estimate the contributions of each signature to each cancer genome.

#### Approach used

- Several mathematical approaches have been used to deconvolute/extract the mutational signatures present in a set of mutational catalogues^3,6,7,9,14–16,39,70–72^. They are all based on the premise that different mutational processes (and thus their signatures) contribute to different extents to different samples within the set.
- Two independently developed methods based on NMF (SigProfiler and SignatureAnalyzer) were applied separately to the sets of mutational catalogues. By using two methods we aimed to provide perspective on the impact different methodologies can have on numbers of signatures generated, signature profiles and attributions. The two methods are described in detail below and the code for both is available (https://www.synapse.org/#!Synapse:syn11801488). Results from the two methods have been compared (https://www.synapse.org/#!Synapse:syn12177006).
- Briefly, SigProfiler employs an elaboration of previously presented approaches for signature extraction and for attribution of mutation counts to mutational signatures in individual tumours^3,4,18,36^.
- Briefly, SignatureAnalyzer employs a Bayesian variant of NMF^6,15,39^. This method enables inferences for the number of signatures through the automatic relevance determination technique and delivers highly interpretable and sparse representations for both signature profiles and attributions at a balance between data fitting and model complexity.
- The methods that SigProfiler and SignatureAnalyzer use for determining the number of extracted signatures are presented in the detailed descriptions of each of these methods, below.
- Both methods assume that the spectra of individual tumours can be represented as linear combinations of signatures. Thus, if the combination of two simultaneously operating mutational processes were to create a signature profile that is not a linear combination of the two, both SigProfiler and SignatureAnalyzer would extract this as a separate signature. We believe this is the case for SBS20, which appears to be due to the simultaneous operation of *POLD1* mutation and mismatch repair deficiency.

#### Role of NMF in extraction and attribution of mutational signatures

- NMF is the approximate representation of a nonnegative matrix *V*, in this case the observed mutational spectra (or profiles) of a set of tumors, as the product of two usually smaller nonnegative matrices, *W* and *H*, which are the signatures and the attributions respectively.
- In our experience, however, calculating a single NMF is rarely sufficient to allow confident extraction and attribution of signatures that reflect the underlying biological mutational processes. There are two main reasons for this:
  - The profiles of extracted signatures can vary substantially depending on the tumour samples present in *V*. For example, this may be especially evident when some tumors in *V* have high numbers of mutations (e.g., samples due to UV exposure or DNA mismatch repair deficiency), while others have low numbers. In situations such as this, signatures due to highly mutagenic processes sometimes capture mutations from other processes and also "bleed" into other signatures.
  - With multiple potentially similar signatures operating, there are multiple possible and reasonably accurate reconstruction solutions for each tumour, often with many small and/or biologically implausible contributions.
- To address these challenges two key additional analytic features have been incorporated into our analyses:
  - Both SigProfiler and SignatureAnalyzer carried out multiple NMFs on different subsets of tumours for signature extraction, and indeed, each signature extraction by SigProfiler entails 1024 NMFs with different random initial conditions. We describe below how we selected representative mutational signature profiles.
  - Both SigProfiler and SignatureAnalyzer developed a process of attributing signature activities to tumours that is separate from the process of extracting (discovering) the signatures.
- The use of multiple extractions to support confidence in results:
  - SignatureAnalyzer, carried out the main extraction procedure on (1) the majority of the PCAWG tumours excluding certain highly mutated tumours and (2) the melanomas, microsatellite-instable tumours, and a single temozolomide-exposed tumour (https://www.synapse.org/#!Synapse:syn11738314).
  - SigProfiler extracted signatures from
    - Separate extraction of SBS, DBS, and ID signatures from all PCAWG whole-genomes together (the main source of the reference mutational signature).
    - Separate extraction of SBS, DBS, and ID signatures from PCAWG whole-genomes with each tumour type examined by itself.
    - Extraction of SBS signatures from all non-PCAWG whole-genomes together.
    - Extraction of SBS signatures from non-PCAWG whole-genomes with each tumour type examined by itself.
    - Separate extraction of SBS and ID signatures from all TCGA exomes together.
    - Separate extraction of SBS and ID signatures from TCGA exomes with each tumour type examined by itself.
    - Separate extraction of SBS and ID signatures from all non-TCGA exomes together.
    - Separate extraction of SBS and ID signatures from non-TCGA exomes with each tumour type examined by itself.
    This allowed the extraction of signatures that were not present in the PCAWG tumours (e.g., SBS42, which has been attributed to haloalkane exposure and seen only in whole exome sequencing data). It also served as an important validation, as extraction of similar signatures from single tumour types and other sample sets supports the correctness of the signature extracted from the PCAWG samples (https://www.synapse.org/#!Synapse:syn12016215).
  - Signature extraction from each tumour type (or from some other subset of cancers) separately has the advantages of:
    - Usually including fewer (and different) mutational signatures in each tumour type sample set than in the set of all cancers together and thus fewer (and different) opportunities for inter-signature interference.
    - Allowing multiple independent opportunities for extraction of a signature that is present in multiple tumour types, and thus of obtaining validation/confirmation of the signature’s existence and profile.
    - Allowing extraction of a signature that may (for a number of reasons) fail to be extracted in analysis of all tumour types together.
    - Providing primary evidence for the existence of the signature in each tumour type.
    - Allowing separation of highly mutated cancer types/samples from cancer types/samples with low mutation burdens.
  - Signature extraction from multiple tumour types together has the advantages of:
    - Usually including more samples with a particular signature than in each individual cancer type and thus being better powered to separate a signature from other partially correlated signatures and/or from signatures with similar profiles.
    - Providing a single profile for a signature rather than the multiple slightly different profiles which emerge from extraction of each tumour type separately.
- The profiles of the mutational signatures extracted from cancer are highly variable. They range from some that have contributions from mutations of all subtypes in the mutation classification (“flat” or “featureless” signatures, e.g., SBS5 and SBS40) to others that are essentially defined by mutations at only one (or a small number) of the mutation subtypes (e.g., signatures SBS2, SBS13, SBS10a and SBS10b). There appears to be less concordance between the results of SigProfiler and SignatureAnalyzer for flat signatures than for signatures with distinct features indicating that generally, these may be more difficult to accurately extract and distinguish from each other. However, there is experimental support for the existence of SBS5 and SBS3^61,68^.
- We represented each signature as a single reference. This selection of a single reference signature does not exclude the possibility that signature profiles may show nuances and further complexity and may vary in different contexts (e.g., in different tissues). The rationale for selecting a single reference signature was the view that this would be a level of granularity useful to most researchers. For those with specialised interests in particular mutational processes and their components, we also provided the signatures extracted from individual tumour types, comprising PCAWG and non-PCAWG genomes and exomes (https://www.synapse.org/#!Synapse:syn12025142).
- Attribution of signatures to cancer samples:
  - The reference signatures from SigProfiler and SignatureAnalyzer were used to estimate the number of mutations due to each signature in each tumour (https://www.synapse.org/#!Synapse:syn11804065).
  - SigProfiler and SignatureAnalyzer differ in their approaches for attributing signatures. However, both incorporate a set of rules based on prior knowledge and biological plausibility, and incorporate techniques to encourage sparsity in the number of signatures attributed to a given tumour.
  - Sparsity (limiting the numbers of signatures and limiting the numbers of signatures attributed to each cancer sample) is an important concept and feature of both SigProfiler and SignatureAnalyzer (both in signature extraction and attribution). Our prior beliefs are that *(i)* there is a limited set of significantly contributing mutational processes (and hence a limited set of mutational signatures) operating to generate somatic mutations across all cancers and *(ii)* that a limited set of mutational processes contribute to individual cancer genomes (as opposed to all mutational signatures contributing to all samples). Our aim in discovering mutational signatures is to reflect the underlying biological processes and to attribute them appropriately. It is not a mathematical exercise in which the main objective and priority is to minimize the difference between *W × H* and the original spectra in *V*. Indeed, if the latter was the main aim, for 96 mutation classes a set of 96 signatures each constituted entirely of mutations in just one class (and therefore ignoring sparsity), will always provide error free reconstruction but will provide absolutely no information about underlying mutational processes.

#### Presentation of the results of signature extraction and attribution from SigProfiler and SignatureAnalyzer

- The results (signatures and attributions) of the two methods have been presented separately. We have done this in preference to combining them. We have handled the two outputs in this way because we believe that this provides a simpler conceptual and technical basis on which the research community can understand the results, can employ the methods in future and can compare results with those shown in this paper. We also do not have a basis for believing that a combined/averaged/overlapping single result set is a better representation of the natural truth than either of the two result sets individually and do not have a well-founded and simple technical approach for combining them. We have, however, provided comparisons of the outputs.
- For brevity and for continuity with previous publications, the results from SigProfiler, a further elaborated version of previously described approaches^3,4,18,36^ that generated the 30 signatures previously shown in COSMICv2^37^, are shown in the main manuscript, and the results from SignatureAnalyzer in supplementary data (https://www.synapse.org/#!Synapse:syn11738307).
- Nomenclature of signatures is based on and extends the nomenclature previously used in COSMIC (COSMICv2, https://cancer.sanger.ac.uk/cosmic/signatures_v2)^37^.
- Both methods analysed each mutation type (SBSs, DBSs and IDs) separately and also together as a composite signature. In future, however, SigProfiler will usually use the separately extracted single base substitution, indel and doublet base substitution signatures as its standard. This generally facilitates portability, and comparison of signature profiles with those from a variety of sample sets including targeted sequences, exomes etc.
- SBS signatures reported in Supplementary Data include possible artefacts (https://cancer.sanger.ac.uk/cosmic/signatures/SBS/ and see below).

#### Quality control: annotating signatures as likely real or a possible artefact

- Sequencing artefacts and differences in analysis pipelines can also generate mutational signatures. We have annotated which signatures are likely real or “possible artefact”.
- There are multiple reasons for believing a signature reflects a biological mutational signature rather than an artefact.
  - The input data supporting the signature seem correct: key mutational features of the putative signature look real in a mapped-read browser such as Integrative Genomics Viewer (IGV, https://software.broadinstitute.org/software/igv/), or characteristic mutations are experimentally confirmed in the tumour and normal samples. Inspection in a mapped read browser is especially important in checking for possible problems in potentially new signatures arising in datasets other than the highly scrutinized and checked PCAWG and TCGA sets. Features associated with experimental, mapping, or other computational artefacts include strong preference for the first read, very low variant allele fractions, variants in regions of low germ-line sequencing coverage, variants found near indels in low-complexity regions, variants from a signature only found in one sequencing centre etc.
  - The 96-mutation profile and additional features (e.g., strand asymmetry, association with replication timing), are known to result from a particular process in experimental systems. Examples: UV, polymerase epsilon proofreading deficiency, aristolochic acid and cisplatin exposure.
  - The putative signature is broadly consistent with previous biochemical knowledge of mutational processes (e.g., preference for G adducts in aflatoxin).
  - The putative signature dominates the spectra of some tumours (column J of https://www.synapse.org/#!Synapse:syn12016215).
  - The putative mutational signature is consistently deciphered from multiple independent datasets; this indicates that the signatures is either a common sequencing artefact or something real.
  - The putative signature correlates with known or suspected mutational exposures, endogenous processes, or repair defects, especially if some of those exposures/processes/repair defects result in overwhelming mutational spectra. Examples: melanoma / fair skin / UV exposure, POLE mutations, MMR deficiency and APOBEC germ line variants.
  - The putative signature correlates with other clinical characteristics, such as age at diagnosis (examples SBS1 and SBS5) or tobacco smoking (SBS4).
  - The mutational signature exhibits a strong transcriptional strand bias; it is hard to imagine an artefact with transcriptional strand bias.
  - The putative signature shows association with other genomic features, such as microindels in homopolymers, replication strand, replication timing, or nucleosome occupancy.

#### Cancer sample sets on which different analyses have been conducted

- Because PCAWG genomes are of high quality with respect to the calling of all mutation types, all our analyses (all types of signature extraction and all types of signature attribution) have been conducted on the 2,780 PCAWG genomes.
- SigProfiler also extracted SBS signatures from the non-PCAWG whole genomes, TCGA exomes, and non-TCGA exomes and attributed SBS signatures to them.
- ID signatures have been extracted and attributed to PCAWG genomes and to a subset of TCGA exomes with large numbers of indels (the latter SigProfiler only). We have not done this for indels in non-PCAWG whole genome sequences and non-TCGA exomes *(i)* because of the unknown and variable accuracy and standardisation of indel mutation calls from different groups generating the data, *(ii)* because in some cases no indel calls were provided by the data generator and *(iii)* because for exomes in most cases there would be very few mutations.
- DBS signatures have been extracted and attributed to PCAWG genomes only. We have not done this for the other categories of samples because of the unknown and variable quality of the mutation calls, the possibility that filters introduced for quality control might deliberately exclude doublet mutations, and the small numbers of doublet mutations in exomes.
- Consistent with the above, composite mutational signatures have only been extracted and attributed for PCAWG genomes.

#### Splitting of mutational signatures

- Certain previously existing single signatures have split into multiple constituent signatures in this analysis. This is likely due to the existence of multiple, partially correlated mutational processes with the same initiating factor (for example, UV exposure) but subsequent differences in underlying mechanisms which differ in intensity in different tissues or other contexts. A previous example of this for which we have allocated different signature numbers is the split of the usually co-occurring but independently varying consequences of APOBEC mutagenesis into signatures SBS2 and SBS13 (https://cancer.sanger.ac.uk/cosmic/signatures/SBS/).
- Depending on the extent of correlation of the two signatures, and the available dataset/statistical power such signatures may manifest as a single signature, overlapping partially separated signatures or as two separate signatures.
- We are aware that splitting of signatures can also be a mathematical artefact. However, we have used multiple extractions to confirm and validate signature splits and applied the principle of sparsity to limit artefactual splits (https://cancer.sanger.ac.uk/cosmic/signatures/SBS/).

#### Better separation compared to COSMICv2 signatures

As described in the manuscript, all mutational signatures previously reported on COSMIC were confirmed in the new set of analyses with median cosine similarity of 0.95. However, the separation between the COSMICv2 mutational signatures (https://cancer.sanger.ac.uk/cosmic/signatures_v2) is much worse compared to the separation between the PCAWG mutational signatures. One can easily discern this by visual examination of signature profiles. For example, in COSMICv2, signatures 5 and 16 have a cosine similarity of 0.90, thus making them hard to distinguish from one another. In contrast, in the current PCAWG analysis, SBS5 and SBS16 have a cosine similarity of 0.65. This allows unambiguously assigning SBS5 and SBS16 to different samples. In the PCAWG analysis, the larger number of samples has allowed reducing the bleeding between signatures and has given more unique and easily distinguishable signatures. One can evaluate the overall separation of a set of mutational signatures by examining the distribution of cosine similarities between the signatures in the set. The COSMICv2 signatures have a median cosine similarity between the signatures in COSMICv2 of 0.238. In contrast, the PCAWG signatures have a much lower median cosine similarity between the signatures in PCAWG of 0.098. This 2-fold reduction in similarity is highly statistically significant (p-value: 9.1 × 10^−25^) and indicates a better separation between the signatures in the current PCAWG analysis.

### Correlations of mutational signature activity with age

Prior to evaluating the association between age and the activity of a mutational signatures, all outliers for both age and numbers of mutations attributed to a signature in a cancer type were removed from the data. Outlier was defined as any value outside three standard deviations from the mean value. A robust linear regression model that estimates the slope of the line and whether this slope is significantly different from zero (F-test; p-value<0.05) was performed using the MATLAB function robustfit (https://www.mathworks.com/help/stats/robustfit.html) with default parameters. The p-values yielded from the F-tests were corrected using the Benjamini-Hochberg procedure for false discovery rate. Results are at https://www.synapse.org/#!Synapse:syn12030687 and https://www.synapse.org/#!Synapse:syn20317940.

### SigProfiler overview

SigProfiler incorporates two distinct steps for identification of mutational signatures based on the previously described methodology^3,4,18,36^. The first step, SigProfilerExtraction, encompasses a hierarchical *de novo* extraction of mutational signatures based on somatic mutations and their immediate sequence context, while the second step, SigProfilerAttribution, focuses on accurately estimating the number of somatic mutations associated with each extracted mutational signature in each sample.

### SigProfilerExtraction

(Note: This phase is termed SigProfiler in the MATLAB code and SigProfilerExtractor in Python). The hierarchical *de novo* extraction approach is an extension of our previous framework for analysis of mutational signatures (Extended Data Figure 8a)^3,18^. Briefly, for a given set of mutational catalogues, the previously developed algorithm was hierarchically applied to an input matrix 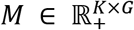 of non-negative integers with dimension *K* × *G*, where *K* is the number of mutation types and *G* is the number of samples. This previously described algorithm deciphers a minimal set of mutational signatures that optimally explains the proportion of each mutation type and estimates the contribution of each signature to each sample. The algorithm uses multiple NMFs to identify the matrix of mutational signatures, 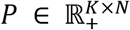, and the matrix of the activities of these signatures, 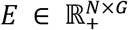, as previously described^3^. The unknown number of signatures, *N*, is determined by human assessment of the stability and accuracy of solutions for a range of values for *N*, as described^3^. The identification of *M* and *P* is done by minimizing the generalized Kullback-Leibler divergence:

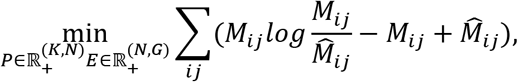

where 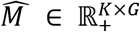 is the unnormalized approximation of *M*, *i.e.*, 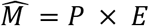. The framework is applied hierarchically to increase its ability to find mutational signatures generating few mutations or present in few samples. In detail, after application to a matrix *M* containing the original samples, the accuracy of reconstructing the mutational spectrum of each sample with the extracted mutational signatures is evaluated. Samples that are well-reconstructed are removed, after which the framework is applied to the remaining sub-matrix of *M*.

Transcriptional strand bias associated with mutational signatures was assessed by applying SigProfilerExtraction to catalogues of in-transcript mutations that capture strand information (192 mutations classes, https://www.synapse.org/#!Synapse:syn12026195). These 192-class signatures were collapsed to strand-invariant 96-class signatures and compared to the signatures extracted from the 96-class data, revealing very high cosine similarities (median 0.90, column F in https://www.synapse.org/#!Synapse:syn12016215).

### SigProfilerAttribution (single sample attribution)

(Note: This phase is termed SigProfilerSingleSample in both the MATLAB and Python code). After signatures are discovered by SigProfilerExtraction, another procedure, SigProfilerAttribution, estimates their contributions to individual samples. For each examined sample, 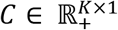, the estimation algorithm involves finding the minimum of the Frobenius norm of a constrained function (see below for constraints) for a set of vectors *S*_*i*=1..*q*_ ∈ Q, where Q is a (not necessarily proper) subset of the set of mutational signatures, P, ie, Q ⊆ P.

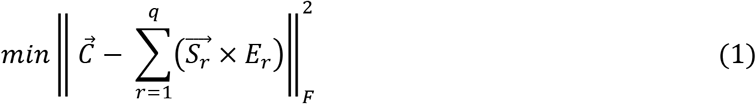

In equation (1), 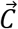 and each 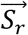 are vectors of *K* nonnegative components reflecting, respectively, the mutational spectrum of a sample and the *r-th* reference mutational signature. All mutational signatures, 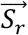, were identified in the SigProfilerExtraction step. Each *E*_*r*_ is unknown scalar reflecting the number of mutations contributed by signature 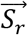 in the mutational spectrum 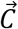. The minimization of equation (1) is always performed under two additional constraints: (i) *E*_*r*_ ≥ 0 and (ii) 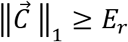; The constrained minimization of equation (1) is performed using a nonlinear convex optimization programming solver using the interior-point algorithm^73^.

SigProfilerAttribution follows a multistep process, wherein equation (1) is minimized multiple times with additional constraints (Extended Data Figure 8b).

In the first phase, the subset *Q* contains all signatures that were found by *SigProfilerExtraction* in the same cancer type as the examined sample. Furthermore, signatures violating biologically meaningful constraints based on transcriptional strand bias and/or total number of somatic mutations are excluded from the set *Q* (https://www.synapse.org/#!Synapse:syn12177009). Further, any 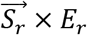 for which the cosine similarity between *Ĉ* and 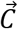 is ≤ 0.01 are sequentially removed, where 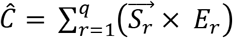. Let *T* be the final set of signatures attributed to the sample at the end of the first phase.

In the second phase, equation (1) is minimized by sequentially allowing each signature, *S*_*r*_ ∈ P\Q,to be added provided that it increases the cosine similarity between *Ĉ* and 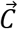 by >0.05. During this second phase, several additional biological conditions are enforced: *(i)* signatures SBS1 and SBS5 are allowed in all samples, *(ii*) if one connected SBS signature is found in a sample than another one is also allowed in the sample (e.g., if SBS17a is found in a sample then SBS17b is allowed in the sample).

### SignatureAnalyzer overview

SignatureAnalyzer employs a Bayesian variant of NMF that infers the number of signatures through the automatic relevance determination technique and delivers highly interpretable and sparse representations for both signature profiles and attributions that strike a balance between data fitting and model complexity. Please see references ^6,15,39^ for more details.

### SignatureAnalyzer signature extraction

In 2,780 PCAWG samples, we applied a two-step signature extraction strategy using 1536 penta-nucleotide contexts for SBSs, 83 ID features, and 78 DBS features. In addition to separate extraction of SBS, ID, and DBS signatures, we performed a "COMPOSITE" signature extraction based on all 1697 features (1536 SBS + 78 DBS + 83 ID). For SBSs, the 1536 SBS COMPOSITE signatures are preferred, and for DBSs and IDs, the separately extracted signatures are preferred.

In step 1 of the two-step extraction process, global signature extraction was performed for the low mutation burden samples (n = 2,624). These excluded hyper-mutated tumours: those with putative polymerase epsilon (POLE) defects or mismatch repair defects (microsatellite instable tumours - MSI), skin tumours (which had intense UV mutagenesis), and one tumour with temozolomide (TMZ) exposure. Because SignatureAnalyzer’s underlying algorithm performs a stochastic search, different runs can produce different results. In step 1 we ran SignatureAnalyzer 10 times and selected the solution with the highest posterior probability. In step 2, additional signatures unique to hyper-mutated samples were extracted (again selecting the highest posterior probability over 10 runs), while allowing all signatures found in the low mutation burden-samples to explain some of the spectra of hyper-mutated samples. This approach was designed to minimize a well-known "signature bleeding" effect or a bias of hyper- or ultra-mutated samples on the signature extraction. In addition, this approach provided information about which signatures are unique to the hyper-mutated samples which is later used when attributing signatures to samples.

### SignatureAnalyzer signature attribution

A similar strategy was used for signature attribution; we performed a separate attribution process for low- and hyper-mutated samples in all COMPOSITE, SBS, DBS, and ID signatures. For downstream analyses, we preferred to use the COMPOSITE attributions for SBSs and the separately calculated attributions for DBSs and IDs. Signature attribution in low-mutation burden samples was performed separately in each tumour type (e.g., Biliary-AdenoCA, Bladder-TCC, Bone-Osteosarc, etc.). Attribution was also performed separately in the combined MSI (n=39), POLE (n=9), skin melanoma (n=107), and TMZ-exposed samples (https://www.synapse.org/#!Synapse:syn11738314). In both groups, signature availability (i.e., which signatures were active or not) was primarily inferred through the automatic relevance determination process applied to the activity matrix H only, while fixing the signature matrix, *W*. The attribution in low-mutation burden samples was performed using only signatures found in the step 1 of the signature extraction. Two additional rules were applied in SBS signature attribution to enforce biological plausibility and minimize a signature bleeding: *(i)* allow signature SBS4 (smoking signature) only in lung and head and neck cases; *(ii)* allow signature SBS11 (TMZ signature) in a single GBM sample. This was enforced by introducing a binary, signature-by-sample, signature indicator matrix Z (1 - allowed and 0 - not allowed), which was multiplied by the *H* matrix in every multiplication update of *H*. No additional rules were applied to ID or DBS signature attributions, except that signatures found in hyper-mutated samples were not allowed in low-mutation burden samples.

### Tests on Synthetic Data

Our goal was to evaluate SignatureAnalyzer (SA) and SigProfiler (SP) on realistic synthetic data. We operationally defined "realistic" as corresponding to either SA’s or SP’s analysis of the PCAWG genome data. SA’s reference signature profiles were based on “COMPOSITE” signatures, consisting of 1536 strand-agnostic single base substitutions (SBSs) in pentanucleotide context, 78 doublet base substitutions and 83 types of small insertions and deletions, for a total of 1,697 mutation types. SP’s reference analysis was based on strand-agnostic single base substitutions in the context of one 5’ and one 3’ base; we term this “SBS96” data. For each test, we generated two sets of "realistic" data: *SP-realistic*, based on SP’s reference signatures and attributions, and *SA-realistic*, based on SA’s reference signatures and attributions, as well as two other types of data that involved using SA profiles with SP attributions and vice versa.

#### Generating synthetic data – overview

For tests (i) through (x) below, Synthetic data for sets of synthetic tumours of a given cancer type, *t*, were generated based on three parameters that were in turn based on the observed statistics for each signature, *s*, in cancer type *t*:

π, the proportion of tumours of cancer type *t* with signature *s*

μ, the mean of log_10_ of the number of *s* mutations across those tumours of type *t* that have signature *s*

σ, the standard deviation of log_10_ of the numbers of *s* mutations across those *t* tumours that have *s*

To generate synthetic data,

(*i*) the proportion of tumours affected by *s* was drawn from the binomial distribution based on π,

(*ii*) the number of mutations due to *s* in an affected tumour was drawn from a normal distribution based on μ and σ.

The code used to generate the synthetic data and summarize SignatureAnalyzer and SigProfiler results is open-source and freely available as the SynSig package: https://github.com/steverozen/SynSig/tree/v0.2.0.

### Description of each suite of synthetic data sets

#### i. Synthetic pancreatic adenocarcinoma (1,000 spectra)

https://doi.org/10.7303/syn18500212.1

#### ii. 2,700 synthetic whole-genome mutational spectra – 300 spectra from each of 9 cancer types

These spectra consist of 300 synthetic spectra from each of the following cancer types: bladder transitional cell carcinoma, oesophageal adenocarcinoma, breast adenocarcinoma, lung squamous cell carcinoma, renal cell carcinoma, ovarian adenocarcinoma, osteosarcoma, cervical adenocarcinoma, and stomach adenocarcinoma. https://doi.org/10.7303/syn18500213.1

#### iii. Mutational spectra generated from combinations of flat, relatively featureless mutational signatures -- version 1

1000 synthetic tumours comprised of 500 synthetic Kidney-RCCs (high prevalence and mutation load from SBS5 and SBS40 signatures) and 500 synthetic ovarian adenocarcinomas (high prevalence of and mutation load from SBS3). This data set embodies tumours with high prevalence of the main flat signatures, SBS3, SBS5, and SBS40, in a realistic context.

https://doi.org/10.7303/syn18500214.1

#### iv. Mutational spectra generated from combinations of flat, relatively featureless mutational signatures -- version 2

1000 synthetic spectra all constructed entirely from SBS3, SBS5, and SBS40, using mutational loads modelled on kidney-RCC (SBS5 and SBS40) and ovarian adenocarcinoma (SBS3). Most synthetic spectra have contributions from all three signatures.

https://doi.org/10.7303/syn18500215.1

#### v. Mutational spectra generated from signatures with overlapping and potentially interfering profiles - version 1

500 synthetic bladder transitional cell carcinomas (high in SBS2 and SBS13) and 500 synthetic skin melanomas (high in SBS7a,b,c,d). The potential interference is between SBS2 (mainly C > T) and SBS7a,b (mainly C > T). https://doi.org/10.7303/syn18500217.1

#### vi. Mutational spectra generated from signatures with overlapping and potentially interfering profiles - version 2

1000 synthetic tumours composed from SBS2 and SBS7a,b. Mutational load distributions were drawn from bladder transitional cell carcinoma (SBS2) and skin melanoma (SBS7a,b). Most spectra contain both signatures. The potential interference is between SBS2 (mainly C > T) and SBS7a,b (mainly C > T). https://doi.org/10.7303/syn18500216.1

#### vii. Mutational spectra generated from combinations of signatures conferring high and low mutation burdens

Based on 500 synthetic non-hypermutated tumours (parameters for SBS1 and SBS5 estimated from colorectal and uterine adenocarcinomas) and 500 hyper-mutated tumours (parameters for SBS26 and SBS44 estimated from hypermutated colorectal and uterine adenocarcinomas). High and low mutation burden tumours are segregated for SignatureAnalyzer (which analyses low mutation burden tumours first, then high-burden tumours). SigProfiler analyses all tumours together.

https://doi.org/10.7303/syn18500218.1

https://doi.org/10.7303/syn18500219.1

https://doi.org/10.7303/syn18500216.1

#### viii. A set of 30 random 96-feature mutational signature profiles and a set of 30 random 1697-feature signature profiles (mimicking COMPOSITE signatures, which have 1697 mutation types)

Each of these are used in two types of exposures, one with more (mean ~15.6) signatures per tumour and one with fewer (mean ~4) signatures per tumour.

https://doi.org/10.7303/syn18500221.1

#### ix. 2,700 whole-exome mutational spectra consisting of 300 synthetic spectra from each of 9 different cancer types

This test data set was generated from ***test ii*** by reducing the number of mutations of each type by 0.013 (approximately ratio of mutation counts between whole exome and whole genome mutational spectra). https://doi.org/10.7303/syn18909829.4

##### Summary of findings

Both SA and SP extracted substantially fewer signatures in this test than in ***test ii***. In particular:

**SA**: SA extracted only 18 signatures from the SA-realistic whole-exome data in this suite, compared to the 40 signatures it extracted from the corresponding whole-genome synthetic data in ***test ii*** and compared to the 39 ground-truth signatures in the synthetic spectra. The average cosine similarity between ground-truth and extracted signatures for the synthetic exome data was 0.863, compared to 0.968 for the signatures extracted from the whole-genome spectra in ***test ii***.

**SP**: SP extracted only 8 signatures from the SP-realistic whole-exome data in this suite, compared to the 19 it extracted from the whole-genome data in ***test ii*** and the 21 ground-truth signatures in the synthetic spectra. The average cosine similarity between ground-truth and extracted signatures for the synthetic exome data was 0.825, compared to 0.965 for the signatures extracted from the whole-genome spectra in ***test ii***.

#### x. 1,350 synthetic whole-genome mutational spectra: 150 spectra from each of 9 cancer types

This test data set consisted of every other tumour from ***test ii***.

##### Summary of findings

On the SA-realistic synthetic data, SA extracted fewer signatures in this data set than in ***test ii***, and in fact the number of signatures extracted was closer to the ground truth and the cosine similarities were there higher. SA over-split in the corresponding set of 2,700 tumours, and we speculate that SA’s tendency to over-split signatures is partly dependent on the number of input spectra, with the result that extraction on 1,350 led to less over-splitting. SP extracted fewer signatures on this data set than on ***test ii***. In particular:

**SA**: SA extracted 38 signatures from the SA-realistic data in this suite, compared to the 40 signatures it extracted from the 2,700 whole-genome spectra in ***test ii*** and compared to the 39 ground-truth signatures. The average cosine similarity between ground-truth and extracted signatures for 1,350 genomes was 0.979 compared to 0.968 for the signatures extracted from the 2,700 whole-genome spectra in ***test ii***.

**SP**: SP extracted 16 signatures from the SP-realistic data in this suite, compared to the 19 signatures it extracted from the 2,700 whole-genome spectra in ***test ii*** and the 21 ground-truth signatures. The average cosine similarity between ground-truth and extracted signatures for the 1,350 spectra was 0.939 compared to 0.965 for the signatures extracted from the 2,700 spectra in ***test ii***.

### xi. Extraction of signatures from exome subsets of PCAWG mutational spectra

Our objective was to further test whether availability of mutations from whole-genome mutational spectra, as opposed to whole-exome spectra, enabled us to extract larger numbers of more accurate mutational signature profiles. In this test, we extracted signatures from mutational spectra that were based on only the exome regions of the actual PCAWG tumours (rather than on the purely synthetic data in ***test ix***). The input data and extraction results are at https://doi.org/10.7303/syn18818766. We next summarize our findings for each of the SBS, DBS, and ID mutational signatures.

#### xi-1 SBS signatures

SignatureAnalyzer on COMPOSITE mutational classes (1536 SBS in pentanucleotide context plus DBS and ID) extracted 12 mutational signature profiles from the whole-exome data, none of which strongly resembled any of the 58 signatures it extracted from the whole-genome data. However, some signatures were unions or splits of the signatures extracted from the whole genome data. For example, WI was a union of the APOBEC signatures BI_COMPOSITE_SBS2_P and BI_COMPOSITE_SBS13_P. More broadly, somewhat recognizable SBS portions of the signatures were combined with the DBS and ID portions of the signatures in difficult-to-interpret combinations. We believe that SBS mutation counts were too low when spread across 1536 mutational classes to support robust mutational signature extraction.

SigProfiler on 96 SBS mutational classes extracted 17 mutational signature profiles from the exome data, compared to 48 that it extracted from the whole-genome data. The median cosine similarity of the exome-extracted signature profiles to the mutational signature profiles extracted from the whole genome data was 0.94. An outlier was SBS-E-2, which was a union of SBS2 and SBS13 (which tend to co-occur).

#### xi-2 DBS signatures

SignatureAnalyzer extracted 2 DBS signatures from the whole-exome data, compared to 15 DBS signatures that it extracted from the full whole genome data. One exome-extracted signature was essentially identical to BI_DBS1 (consisting almost entirely of CC > TT mutations), and one somewhat similar to BI_DBS2 (mostly CC > AA) but with many other mutational classes in addition.

SigProfiler extracted 3 DBS signatures from the whole-exome data, compared to the 11 DBS signatures that it extracted from the whole genome data. The exome-extracted signatures were good approximations of DBS1, DBS2, and DBS10 (cosine similarities 1, 0.93, and 0.98).

#### xi-3 ID signatures

SignatureAnalyzer extracted 4 ID signatures from the whole-exome data, compared to 29 ID signatures extracted from the whole-genome data. It extracted close approximations of BI_ID1_P and BI_ID2_P with cosine similarities 0.97 and 0.94. These are insertions (signature W.3) and deletions (signature W.1) of T:A in poly T:A. SignatureAnalyzer extracted 2 additional signatures. One of these (W.4) was a version of BI_ID4_P with several mutational classes absent. The other (W.2) appeared to be a union of many of the remaining ID signatures.

SigProfiler extracted 6 ID signatures from the whole-exome data, compared to the 17 ID signatures that it extracted from the whole genome data. Signatures ID-E-1, ID-E-2, ID-E-3, and ID-E-4 were good approximations of ID1, ID2, ID3, and ID4, respectively. An additional signature, ID-E-5, was approximately a union of ID6 and ID8. The remaining signature, ID-E-6 was a partial version (deletions in C homopolymers only) of ID7.

### Detailed Summary of Results (including links to input synthetic data sets and the signature profiles extracted)

https://doi.org/10.7303/syn18497223 provides a table with the number of signatures extracted by SigProfiler and SignatureAnalyzer for each synthetic data set and the cosine similarities to the input ground-truth signatures. See above for overall interpretation of the results.

### Data Availability

Data are available at https://www.synapse.org/#!Synapse:syn11726601/wiki/513478. All figures and extended data figures have associated raw data.

### Code Availability

SigProfiler is available both as a MATLAB framework and as a Python package. In both cases, SigProfiler is fully functional, free, and open-source tool distributed under the permissive 2-Clause BSD License. SigProfiler in MATLAB can be downloaded from: https://www.mathworks.com/matlabcentral/fileexchange/38724-sigprofiler

SigProfiler in Python can be downloaded from: https://github.com/AlexandrovLab/SigProfilerExtractor. SignatureAnalyzer code is available at https://www.synapse.org/#!Synapse:syn11801492. The code used to generate the synthetic data and summarize SignatureAnalyzer and SigProfiler results is open-source and freely available as the SynSig package: https://github.com/steverozen/SynSig/tree/v0.2.0 <u>under the GPL3 license</u>.

## Extended Data Figure and Table Legends

**Extended Data Figure 1.**
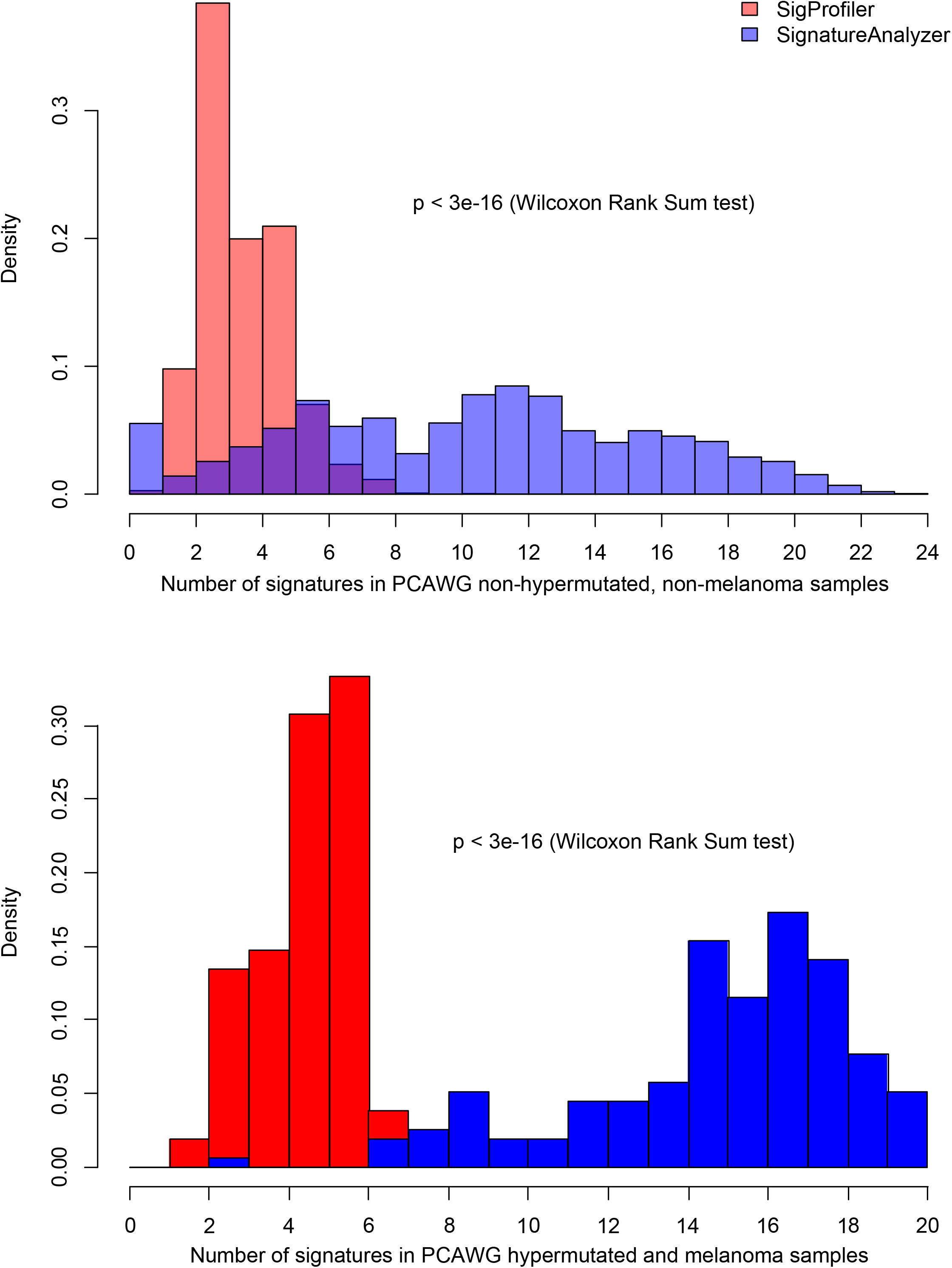
Histogram of number of signatures attributed in each of 2,780 PCAWG samples by SigProfiler and SignatureAnalyzer. Hypermutated tumours and melanomas (156) are listed at https://www.synapse.org/#!Synapse:syn11738314.

**Extended Data Figure 2.**
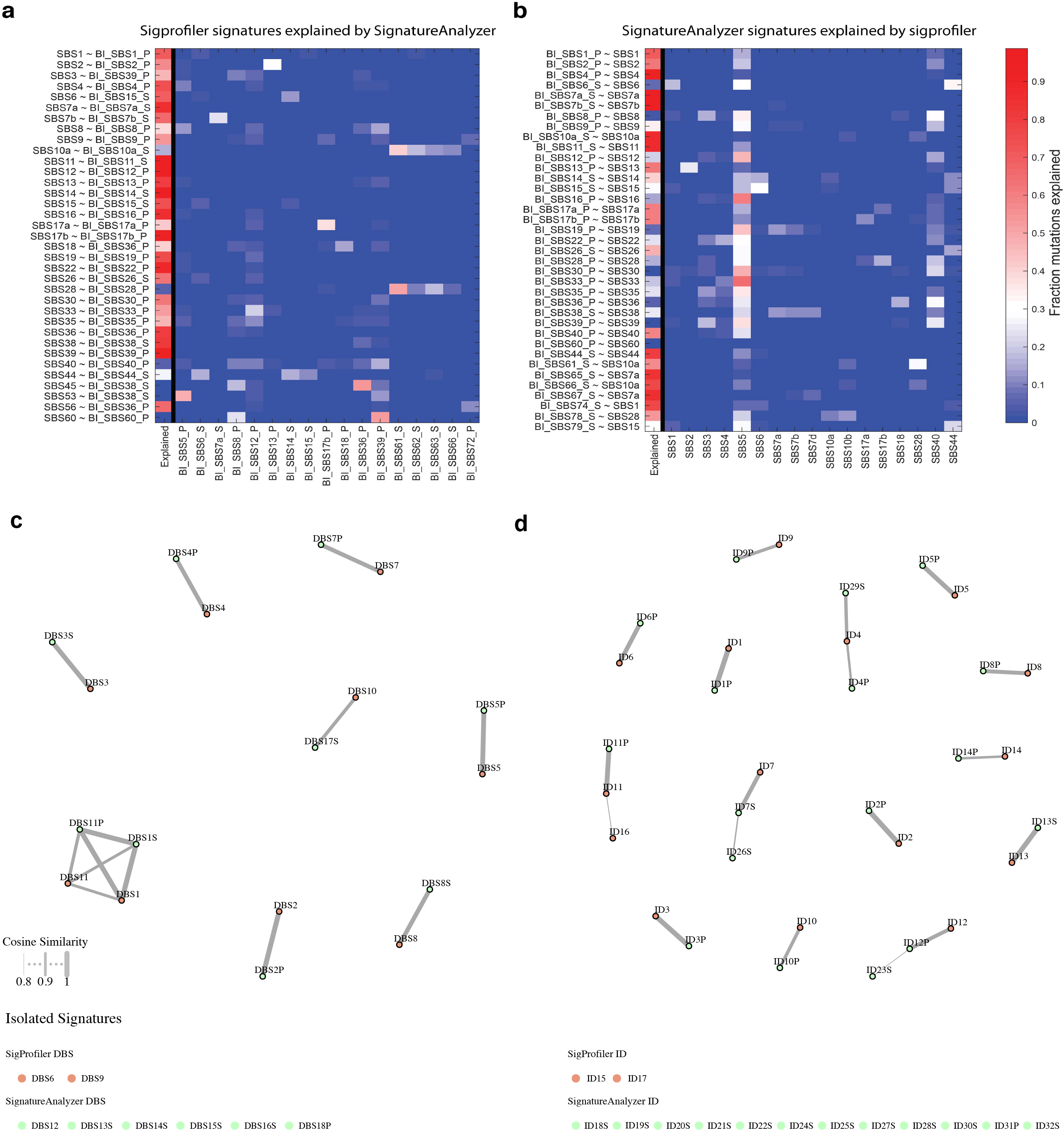
Comparisons between SigProfiler and SignatureAnalyzer results. Comparison of the attributions for corresponding SigProfiler ***(a)*** and SignatureAnalyzer ***(b)*** signatures. Each of the SBS signatures extracted by SigProfiler and SignatureAnalyzer was paired with the signature of highest cosine similarity in the extraction by the other method (if one with >0.85 cosine similarity exists). The first column of the plot corresponds to the fraction of mutations assigned by one method (summed across samples and mutation types) that were also assigned by the other method. The remaining mutations were then re-distributed to the other signatures in the extraction, weighted by their relative probabilities of having been generated by each signature, and the resulting fraction of mutations is plotted. Signatures on the x-axis are only shown if they contribute at least 0.1 fraction of mutations to at least one signature on the y-axis. Cosine similarities between SigProfiler and SignatureAnalyzer DBS ***(c)*** and ID ***(d)*** signatures. Brown nodes represent SigProfiler signatures; green nodes represent SignatureAnalyzer signatures. Matches with cosine similarities > 0.8 are show as edges, with the width of the edge indicate the strength of the similarity. The locations of the nodes have no significance. Signatures with no matches of > 0.8 cosine similarity are show below. Note that SigProfiler ID15 and ID17 were extracted from data that were not analysed by SignatureAnalyzer. Suffixes ‘P’ and ‘S’ on SignatureAnalyzer signature names indicate (1) signatures extracted from non-hypermutated, non-melanoma tumours and (2) hypermutated and melanoma tumours, respectively.

**Extended Data Figure 3.**
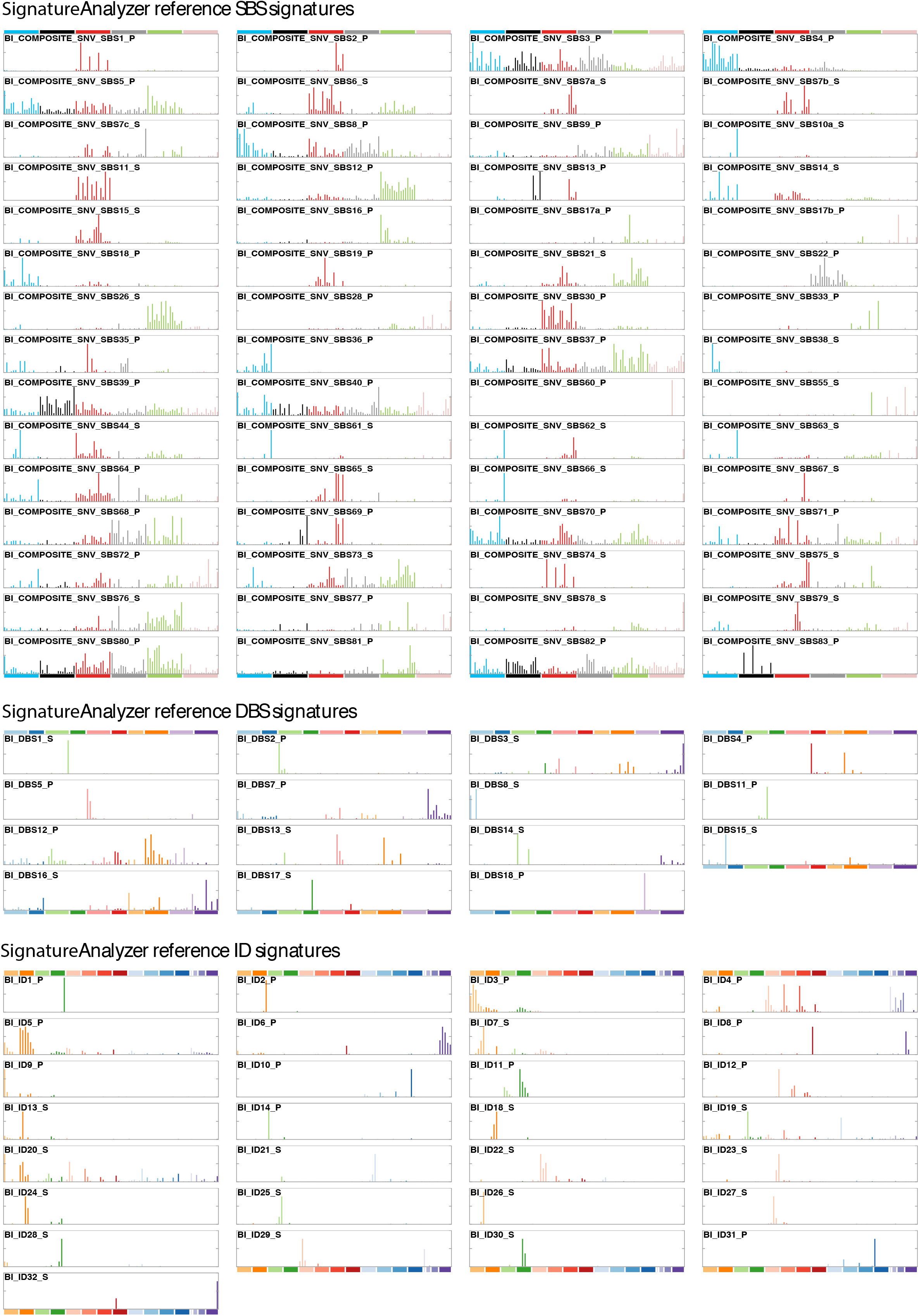
SignatureAnalyzer reference signatures. See legend of main text Figure 2.

**Extended Data Figure 4.**
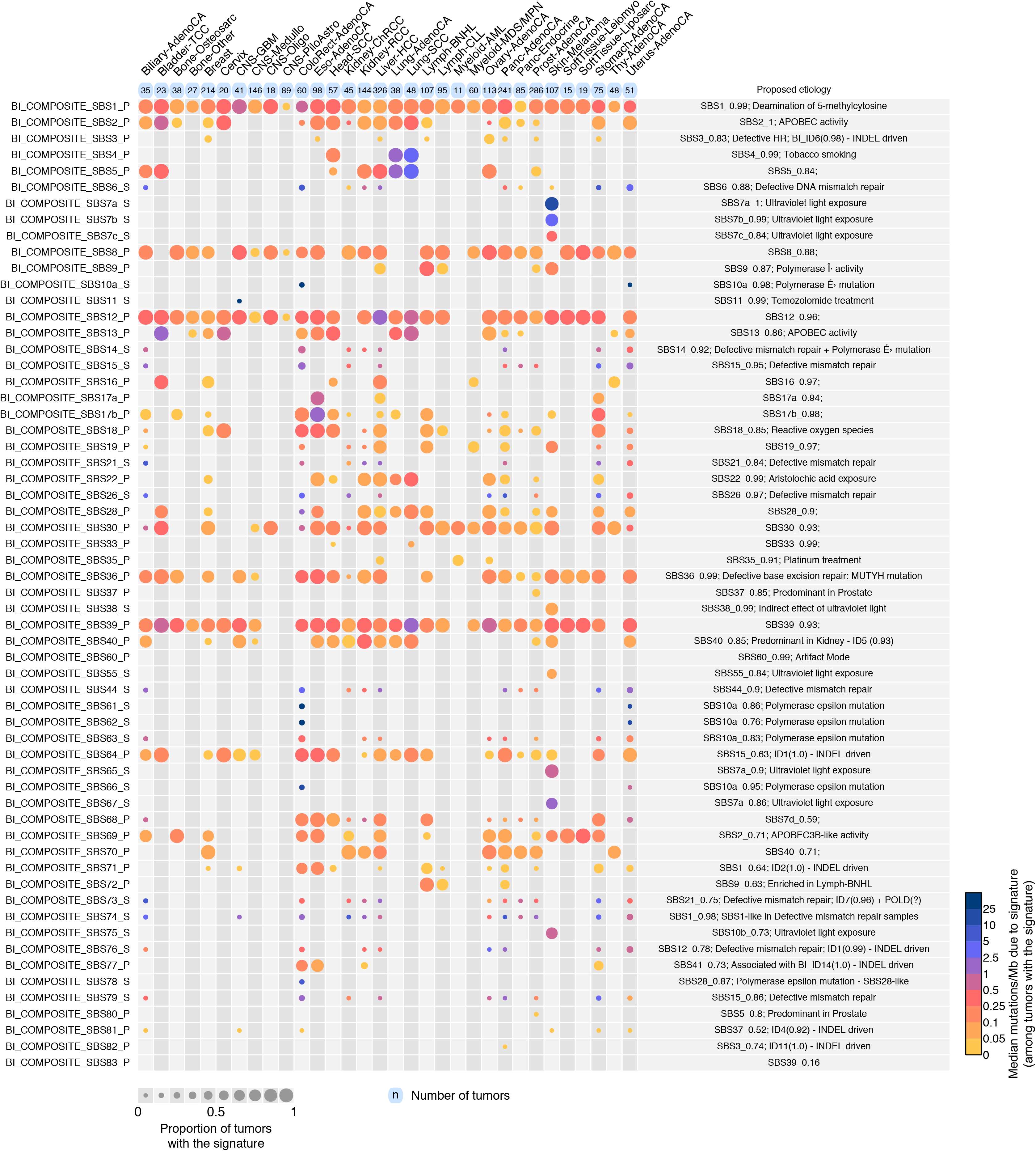
The number of SBS mutations attributed to each mutational signature for each cancer type over the 2,780 PCAWG tumours by SignatureAnalyzer. See main text Figure 3 for explanation.

**Extended Data Figure 5.**
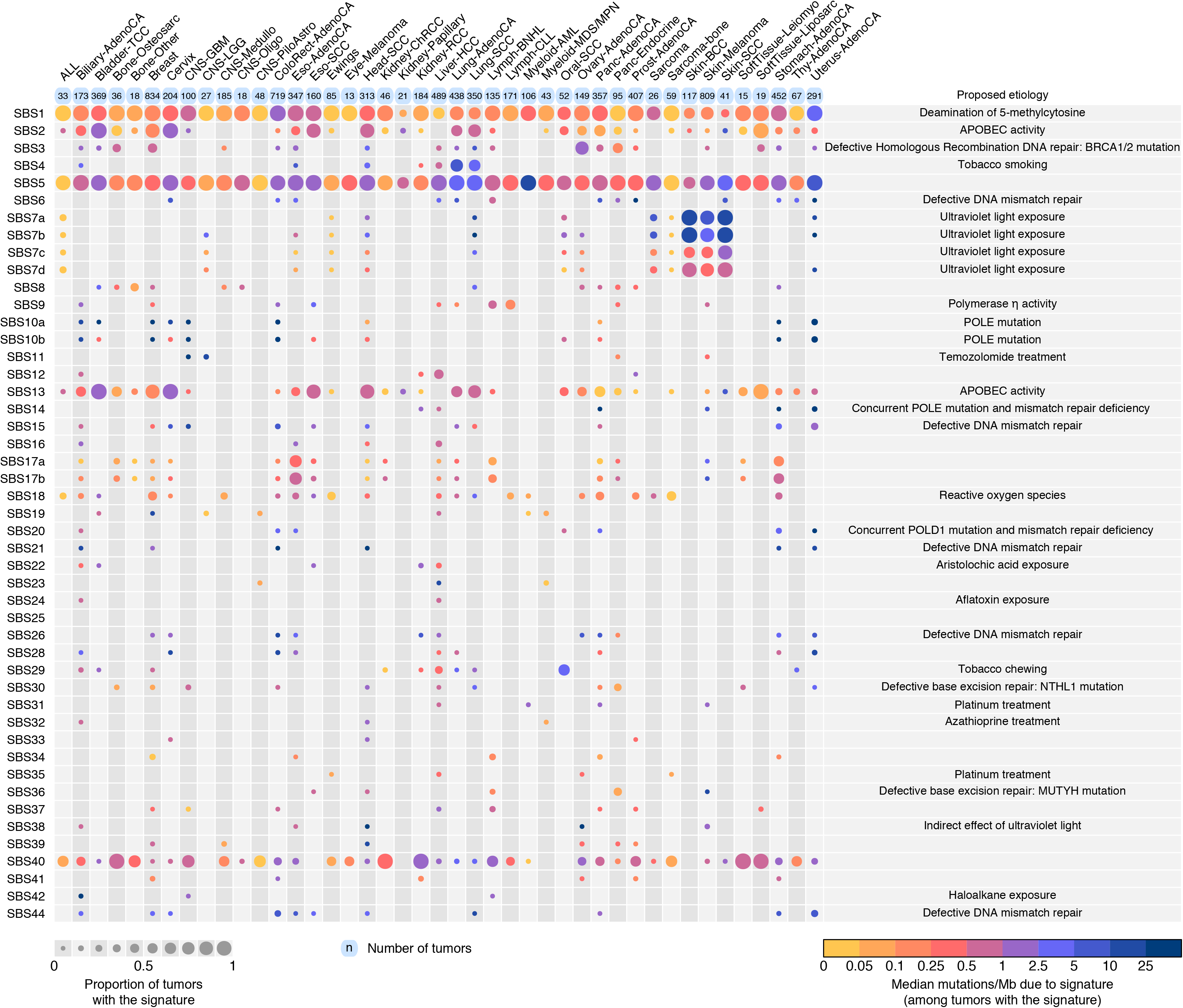
The number of SBS mutations attributed to each mutational signature to each cancer type over the complete set of 23,829 cancer samples analysed by SigProfiler. See main text Figure 3 for explanation.

**Extended Data Figure 6.**
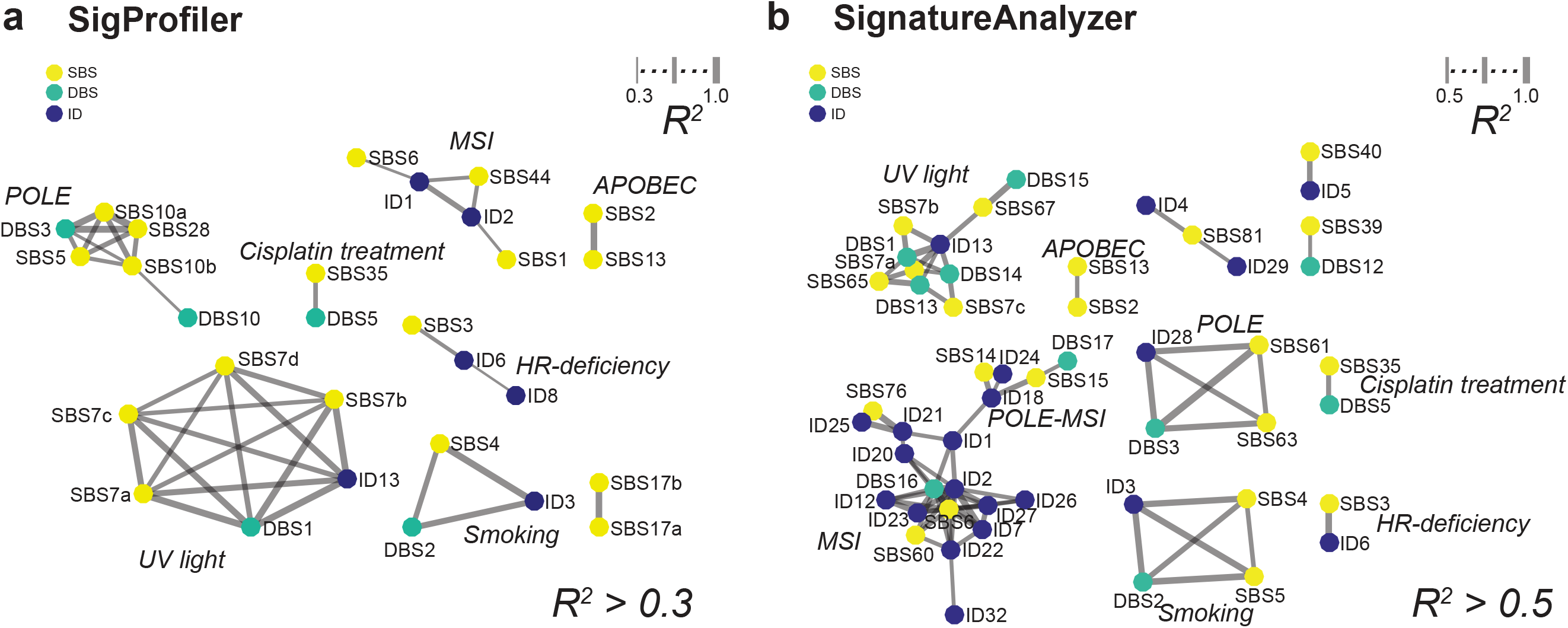
Associations of between SBS, DBS, and ID signature activities for SigProfiler *(a)* and SignatureAnalyzer *(b)*. Each node represents an SBS (light green), DBS (dark green) or ID (black) signature. Any two signatures with sample attributions that significantly correlated with R^2^ > 0.3 (SigProfiler) or > 0.5 (SignatureAnalyzer) are connected by edges. Edge widths are proportional to the strength of the correlation. Signatures with no significant correlation to any other signature above the relevant threshold are not shown. Signature locations are fit for display purposes only and do not indicate similarity.

**Extended Data Figure 7.**
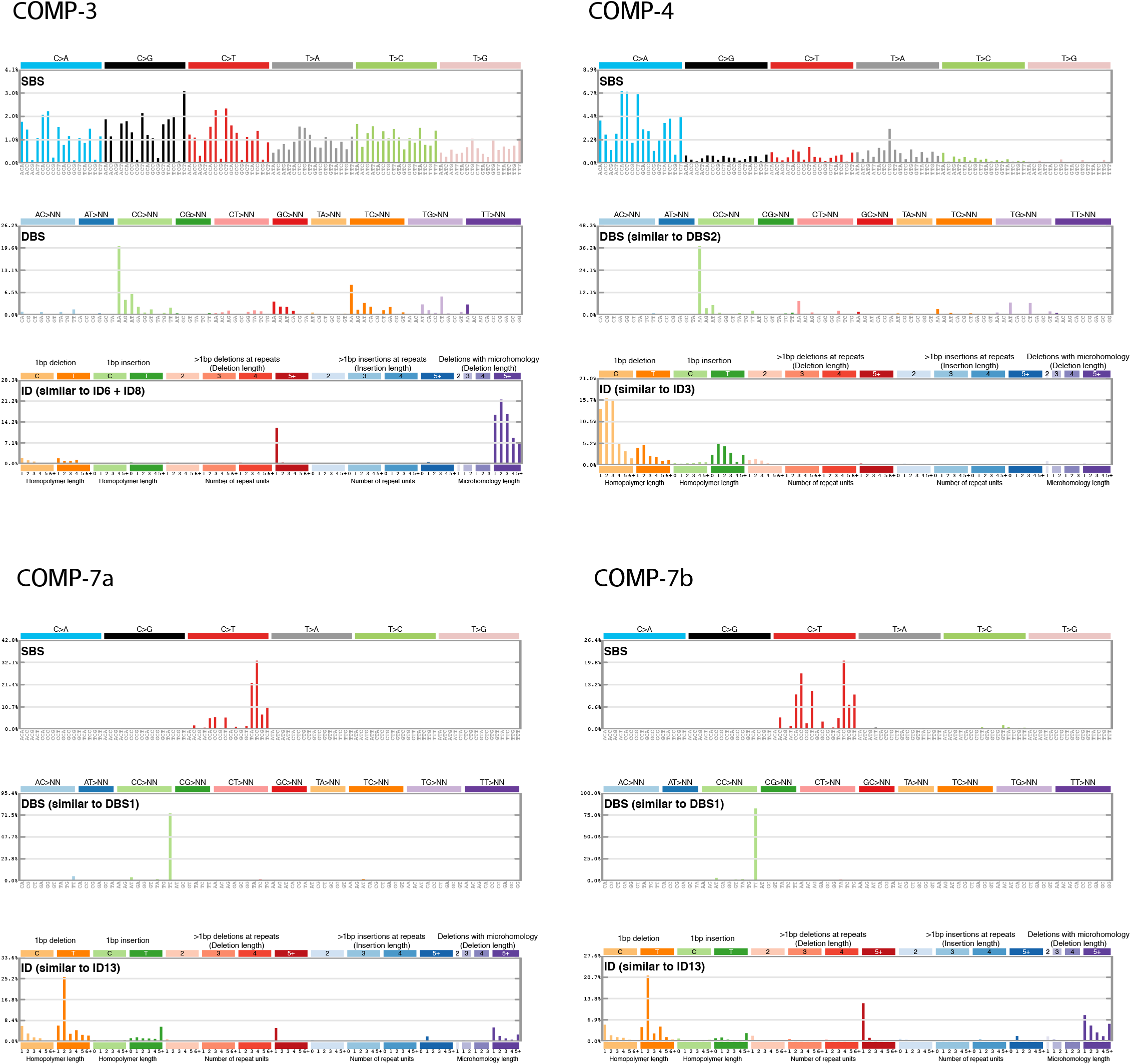
Mutational signatures extracted from the composite feature set consisting of SBSs in pentanucleotide context, DBSs, and IDs. For each of the four composite mutational signatures shown, the top panel is the SBS signature collapsed to 96 SBS classes, the middle panel is the co-extracted DBS signature, and the lower panel is the co-extracted ID signature. Note the similarities between the DBS portion of Composite 4 and DBS2, between the ID portion of Composite 4 and ID3, and other similarities noted in the figure.

**Extended Data Figure 8.**
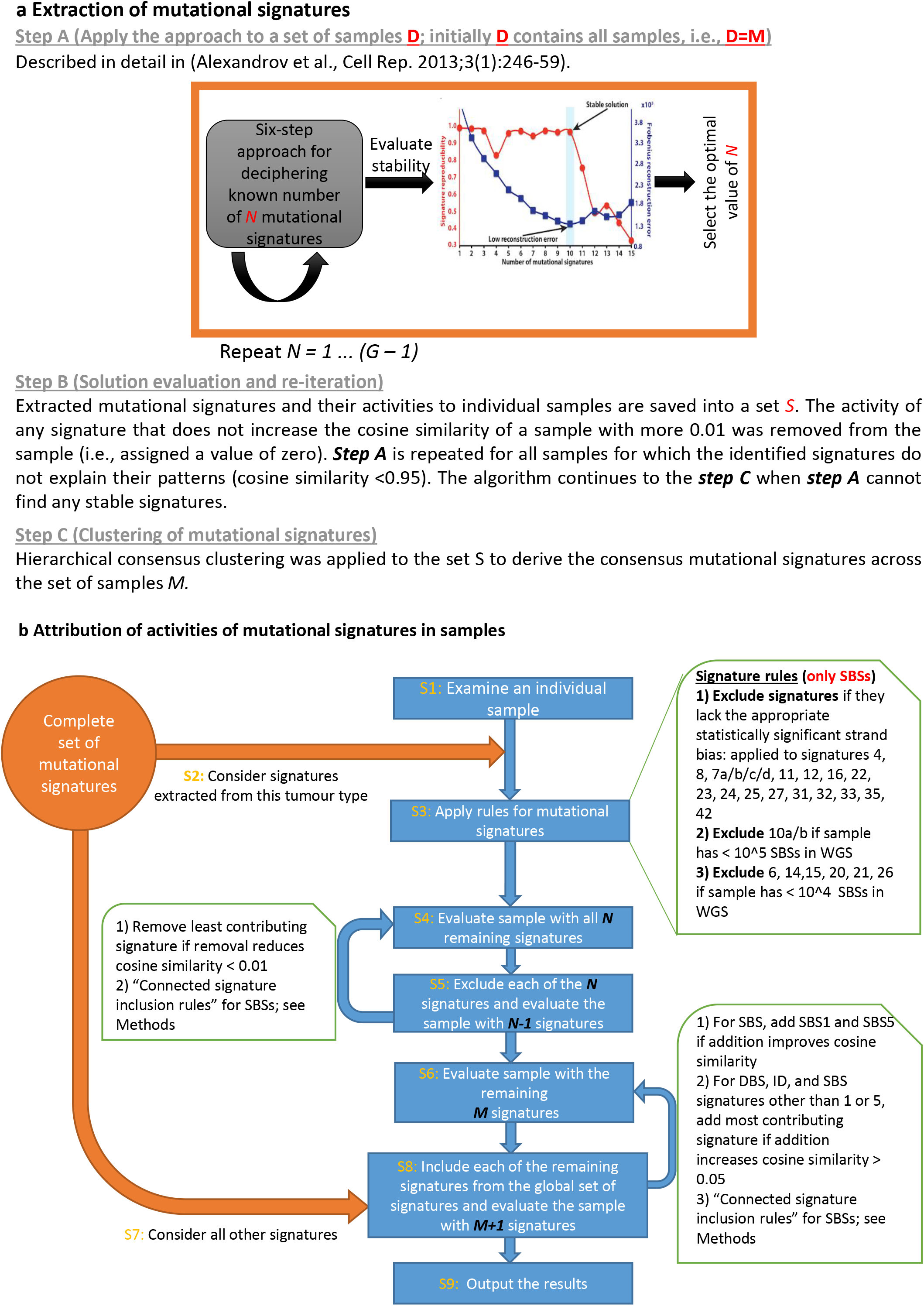
SigProfiler signature extraction *(a)* and attribution *(b)*. See Methods for description.

**Extended Data Table 1.** The number of DBSs is proportional to the number of SBSs with the exception of a few cancer types (ColoRect-AdenoCA, Lung-AdenoCA, Lung-SCC, Skin-Melanoma) analysed by the following linear regression (computed by an R function call): glm(DBS.counts ~ SBS.counts + Cancer.Types).

|  | Estimate | Std.Error | t value | Pr(>\|t\|) |  |
| --- | --- | --- | --- | --- | --- |
| (Intercept) | 5.61E+00 | 8.76E+01 | 0.064 | 0.9489 |  |
| SBS.counts | 3.74E-03 | 1.25E-04 | 29.841 | <2.00E-16 | *** |
| Bladder-TCC | 1.32E+01 | 1.39E+02 | 0.095 | 0.92432 |  |
| Bone-Osteosarc | 2.18E+00 | 1.21E+02 | 0.018 | 0.98567 |  |
| Bone-Other | -2.81E+00 | 1.33E+02 | -0.021 | 0.9831 |  |
| Breast | 5.32E+00 | 9.44E+01 | 0.056 | 0.95511 |  |
| Cervix | -1.06E+01 | 1.45E+02 | -0.073 | 0.94185 |  |
| CNS-GBM | -2.81E+01 | 1.19E+02 | -0.236 | 0.81352 |  |
| CNS-Medullo | -7.04E+00 | 9.75E+01 | -0.072 | 0.94239 |  |
| CNS-Oligo | -1.03E+01 | 1.50E+02 | -0.069 | 0.94539 |  |
| CNS-PiloAstro | -5.87E+00 | 1.03E+02 | -0.057 | 0.95467 |  |
| ColoRect-AdenoCA | -4.11E+02 | 1.12E+02 | -3.667 | 0.00025 | *** |
| Eso-AdenoCA | -1.56E+01 | 1.02E+02 | -0.153 | 0.87838 |  |
| Head-SCC | 5.27E+01 | 1.11E+02 | 0.474 | 0.63541 |  |
| Kidney-ChRCC | -3.14E+00 | 1.17E+02 | -0.027 | 0.97857 |  |
| Kidney-RCC | 5.61E+01 | 9.76E+01 | 0.574 | 0.56584 |  |
| Liver-HCC | 7.82E+01 | 9.21E+01 | 0.849 | 0.39575 |  |
| Lung-AdenoCA | 5.02E+02 | 1.21E+02 | 4.136 | 3.63E-05 | *** |
| Lung-SCC | 5.85E+02 | 1.15E+02 | 5.078 | 4.08E-07 | *** |
| Lymph-BNHL | 1.04E+01 | 1.01E+02 | 0.103 | 0.91765 |  |
| Lymph-CLL | -4.30E+00 | 1.02E+02 | -0.042 | 0.96655 |  |
| Myeloid-AML | -1.89E+00 | 1.79E+02 | -0.011 | 0.99156 |  |
| Myeloid-MDS/MPN | -7.43E+00 | 1.10E+02 | -0.067 | 0.94622 |  |
| Ovary-AdenoCA | 3.59E+01 | 1.00E+02 | 0.358 | 0.72023 |  |
| Panc-AdenoCA | -8.34E-01 | 9.37E+01 | -0.009 | 0.99289 |  |
| Panc-Endocrine | -5.70E+00 | 1.04E+02 | -0.055 | 0.95628 |  |
| Prost-AdenoCA | 2.52E+00 | 9.27E+01 | 0.027 | 0.97831 |  |
| Skin-Melanoma | 1.67E+03 | 1.02E+02 | 16.47 | <2.00E-16 | *** |
| SoftTissue-Leiomyo | 5.98E+00 | 1.60E+02 | 0.037 | 0.97016 |  |
| SoftTissue-Liposarc | 7.77E+00 | 1.48E+02 | 0.053 | 0.95804 |  |
| Stomach-AdenoCA | -3.04E+01 | 1.06E+02 | -0.287 | 0.77417 |  |
| Thy-AdenoCA | -4.80E+00 | 1.15E+02 | -0.042 | 0.96676 |  |
| Uterus-AdenoCA | -1.25E+02 | 1.14E+02 | -1.096 | 0.27304 |  |

**Extended Data Table 2.** Numbers of insertion/deletion mutations due to ID1, ID2, and all other ID signatures in hypermutators and non-hypermutators.

| **Signature** | **Hypermutators** |  | **Non-hypermutators** |  | **All Tumours** |  |
| --- | --- | --- | --- | --- | --- | --- |
|  | **Count** | **Fraction** | **Count** | **Fraction** | **Count** | **Fraction** |
| **ID1** | 593,935 | 0.236 | 399,633 | 0.276 | 993,568 | 0.250 |
| **ID2** | 1,838,867 | 0.730 | 252,893 | 0.174 | 2,091,760 | 0.527 |
| **ID1+ID2** | 2,432,802 | 0.966 | 652,526 | 0.450 | 3,085,328 | 0.777 |
| **Other ID signatures** | 85,038 | 0.034 | 797,964 | 0.550 | 883,002 | 0.223 |
| **Total** | 2,517,840 | 1 | 1,450,490 | 1 | 3,968,330 | 1 |

